# Asymmetric clustering of centrosomes defines the early evolution of tetraploid cells

**DOI:** 10.1101/526731

**Authors:** Nicolaas C. Baudoin, Kimberly Soto, Olga Martin, Joshua M. Nicholson, Jing Chen, Daniela Cimini

## Abstract

Tetraploidy has long been of interest to both cell and cancer biologists, partly because of its documented role in tumorigenesis. A common model proposes that the extra centrosomes that are typically acquired during tetraploidization are responsible for driving tumorigenesis. However, this model is inconsistent with the observation that tetraploid cells evolved in culture lack extra centrosomes. This observation raises questions about how tetraploid cells evolve and more specifically about the mechanisms(s) underlying centrosome loss. Here, using a combination of fixed cell analysis, live cell imaging, and mathematical modeling, we show that populations of newly formed tetraploid cells rapidly evolve *in vitro* to retain a near-tetraploid chromosome number while losing the extra centrosomes gained at the time of tetraploidization. This happens through a process of natural selection in which tetraploid cells that inherit a single centrosome during a bipolar division with asymmetric centrosome clustering are favored for long-term survival.

---

Organismal polyploidy is confined to certain taxa, but many species across the tree of life are thought to have had polyploid ancestors at some point in their evolutionary history^1,2,3,4^ and polyploidy is thought to contribute to speciation and evolution^5,6^.

In vertebrates, organismal polyploidy is rare, and among mammals it has only been described in a single species^7^. However, within individual diploid mammals, some tissues physiologically develop to have a higher ploidy than the majority of somatic cells. Polyploidy can also occur outside of the context of normal development, and is linked with both pathology (particularly cancer^8^) and aging^9^. Tetraploid cells are commonly found in premalignant lesions and tumors at different stages^10,11,12^. Furthermore, meta-analysis of catalogued tumor genomes has provided evidence that close to 40% of all cancers – even those that were not tetraploid at the time of sampling – had a tetraploid intermediate stage at some point during tumor evolution^13^. Consistent with this, several studies have shown a direct, causative link between tetraploidy and tumorigenesis^14,15^.

In proliferating cells, tetraploidy can arise via abnormal cell cycle events, including cytokinesis failure, cell fusion, endoreduplication, and mitotic slippage^12,16,17,18,19^. Importantly, most of these events result in the concomitant acquisition of extra centrosomes along with genome duplication. In cultured cells, extra centrosomes promote chromosomal instability^20,21^ and invasive/migratory behavior^22^. Recent studies have also shown that extra centrosomes promote and in some cases are sufficient to drive tumorigenesis *in vivo*^23,24^.

Based on these studies, it has been speculated that the extra centrosomes emerging as a result of tetraploidization may drive chromosomal instability and, in turn, tumorigenesis^25^. However, anecdotal reports^22,26,27^ have indicated that single cell clones of tetraploid or near-tetraploid cells displayed normal centrosome numbers. This suggests that our understanding of the early evolution of tetraploid cells is very partial and our view of how centrosome and chromosome numbers evolve after tetraploidization needs to be revisited. To address this problem, we studied the time period immediately following cytokinesis failure and investigated how centrosome and chromosome numbers change in newly formed tetraploid cells. Following the observation that centrosome, but not chromosome, number rapidly returns to normal, we combined computational and experimental approaches to identify the specific cellular mechanism underlying the loss of extra centrosomes.

## Results

To investigate the early consequences of tetraploidy and the evolution of newly formed tetraploid cells, we induced cytokinesis failure by dihydrocytochalasin B (DCB) treatment for 20 hr (**Figure 1a**) in both DLD-1 (pseudodiploid colorectal cancer cells) and p53^−/−^ hTERT-immortalized RPE-1 cells^28^ (hereafter referred to as RPE-1 p53^−/−^). Cells generated by this method are referred to, throughout the paper, as ‘newly formed tetraploid cells’ (text) or ‘4N new’ (figures).

**Figure 1.**
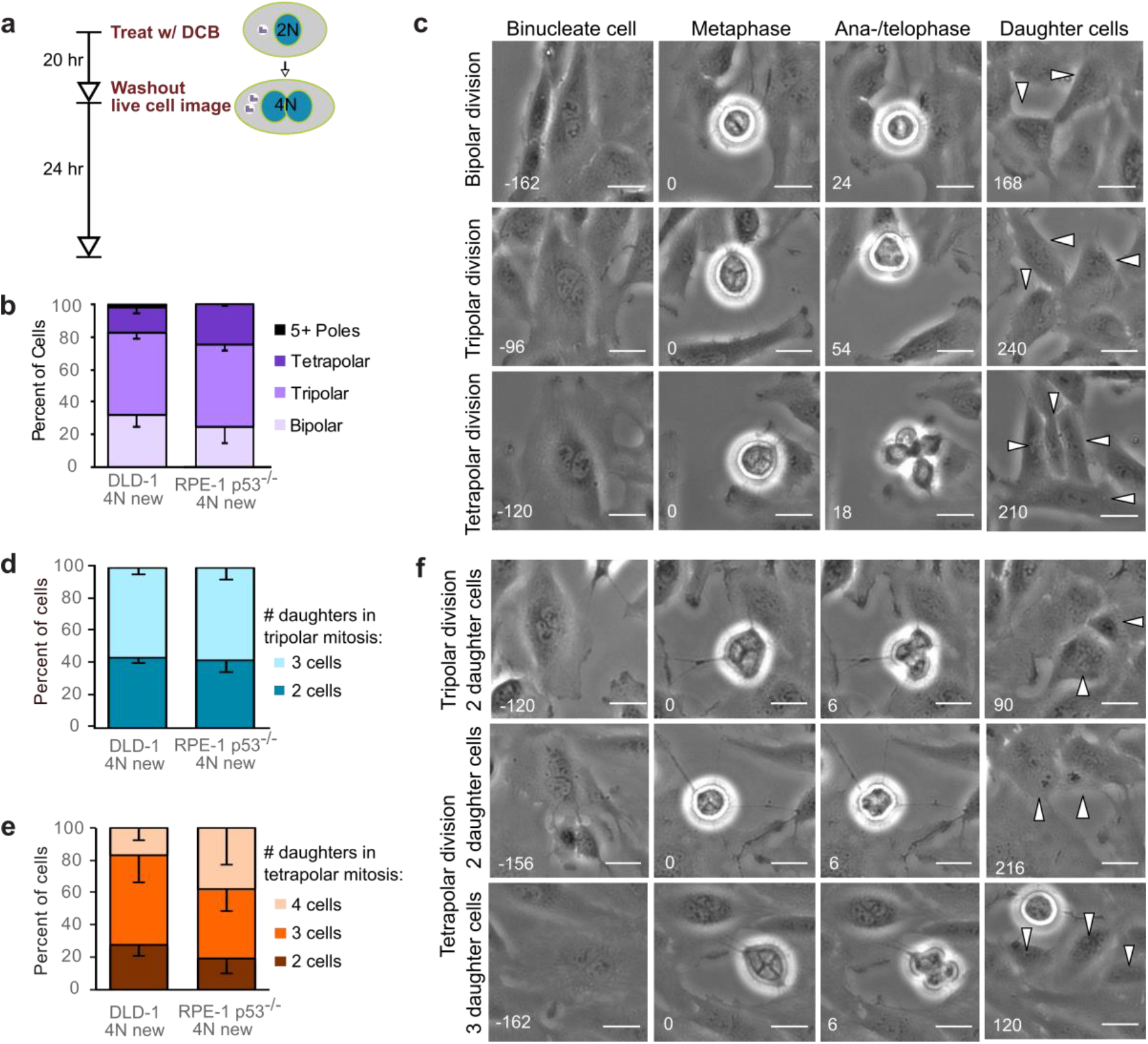
Newly formed tetraploid cells undergo diverse fates in their first mitotic division. (a) Experimental approach for generating newly formed tetraploid cells and performing live cell imaging. (b) Quantification of the type of division observed in the first cell division of newly formed tetraploid cells; characterization was performed at ana-/telophase. (c) Examples of bipolar (top), tripolar (middle), and tetrapolar (bottom) divisions. (d-e) Quantification of incomplete cytokinesis in tripolar (d) and tetrapolar (e) divisions. (f) Examples of multipolar divisions with incomplete cytokinesis such that multiple anaphase poles are clustered into a single daughter cell. Error bars in all graphs represent S.E.M. from three separate imaging sessions. All scale bars, 25 μm. Arrowheads in all images point to a single daughter cell.

### Newly formed tetraploid cells undergo diverse fates in their first mitotic division

To determine the fate of the first tetraploid mitosis, we performed live-cell phase contrast microscopy for 24 hrs following DCB washout (**Figure 1a**; note, newly formed tetraploid cells can easily be identified by the presence of two nuclei). We found that multipolar divisions were frequent in both cell types (**Figure 1b, c**), consistent with the acquisition of extra centrosomes, along with extra chromosomes, upon cytokinesis failure and with the ability of extra centrosomes to promote formation of multipolar mitotic spindles. Only 25-30% of newly formed tetraploid cells underwent bipolar anaphase, indicating that centrosome clustering does not occur efficiently in these newly formed tetraploid cells. However, when we followed these multipolar mitoses through cytokinesis, we found that tripolar anaphases did not always generate three daughter cells and similarly tetrapolar anaphases rarely resulted in four daughter cells (**Figure 1d-f**). Instead, the DNA corresponding to two or more anaphase poles was often enclosed in a single daughter cell, giving rise to binucleated or, rarely, trinucleated daughter cells (**Figure 1d-f**), consistent with previous observations^29,30^. Moreover, we quantified DNA fluorescence at the spindle poles of bipolar and multipolar anaphase cells as an estimate of the proportion of the genome that segregated to each pole (**Figure S1**). This analysis showed that chromosome distribution to the daughter cells was balanced in bipolar divisions (**Figure S1a, c**) but greatly deviated from an equal distribution in multipolar mitoses (**Figure S1b, c**).

### Highly aneuploid karyotypes form early in the evolution of tetraploid cells, but quickly disappear from the population

We next investigated how these early cell divisions after tetraploidization may impact the karyotype of the proliferating cell population. To this end, we carried out a time-course experiment in which we performed chromosome counting in the cell population after the 20 hr DCB treatment and every two days thereafter for a 12-day period (**Figure 2a-b**). Immediately after drug washout, we observed a tetraploid fraction of 90% and 60% of the population for DLD-1 and RPE-1 p53^−/−^ cells, respectively (**Figures 2c, d, day 0**). Two days after DCB washout, we observed high frequencies of cells with karyotypes in the hypotetraploid/hyperdiploid range (**Figures 2b, middle panel; c – e, g**). It is easy to imagine that these highly aneuploid karyotypes may originate from multipolar mitoses. However, the highly abnormal karyotypes rapidly decreased over the course of the experiment and were virtually eliminated from the DLD-1 population by day 12, leaving sub-populations of near-diploid cells (presumably derived from cells that did not respond to the initial DCB treatment) and near-tetraploid cells (**Figures 2b, middle panel; c-e, g**). The appearance and disappearance of highly aneuploid cells corresponded to an increase followed by a decline in the fraction of dead cells between day 2 and 6 (**Figures 2f, h**).

**Figure 2.**
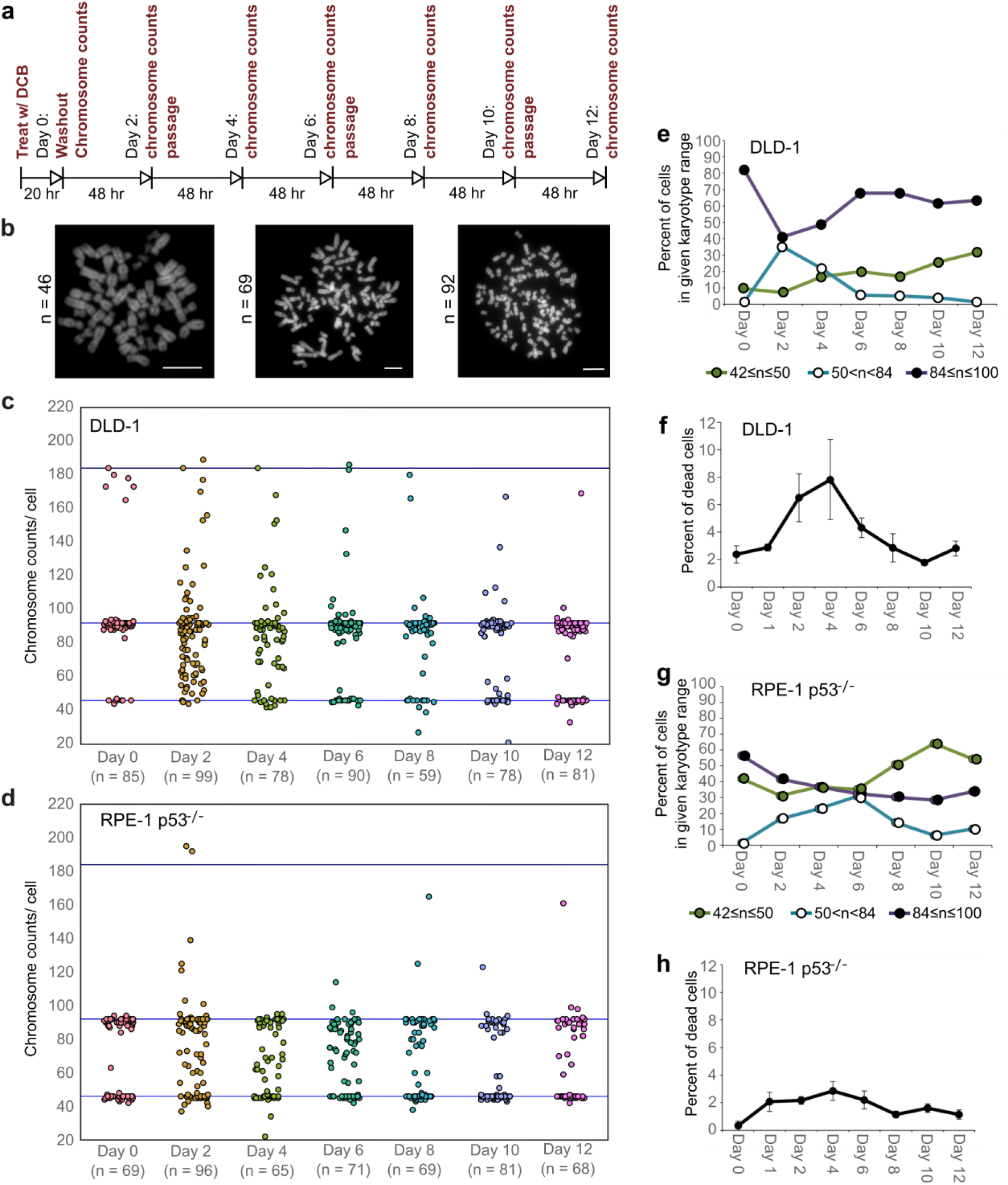
High degrees of aneuploidy appear and rapidly disappear following experimental induction of tetraploidization. (a) Experimental design for time course experiments designed to analyze karyotype evolution in newly formed tetraploid cells over a 12-day time period. (b) Example chromosome spreads from cells with diploid (left), highly aneuploid (middle), and tetraploid (right) karyotypes. Scale bars, 10 μm. (c-d) 12-day time course analysis of chromosome number evolution in newly formed tetraploid DLD-1 (c) and newly formed tetraploid RPE-1 p53^−/−^ (d) cells. (e) Quantification (from the data in c) of the fraction of cells that are near-diploid, highly aneuploid, or near-tetraploid. (f) Time course analysis of cell death in newly formed tetraploid DLD-1 cells. (g) Quantification (from data in d) of the fraction of cells that are near-diploid, highly aneuploid, or near-tetraploid. (h) Time course analysis of cell death in newly formed tetraploid RPE-1 p53^−/−^ cells. Karyotype evolution data were obtained from two biological replicates. Error bars in f and h represent S.E.M. from three biological replicates.

To explore whether the possible karyotypic outcomes of a multipolar division could explain the disappearance of cells with highly aneuploid karyotypes, we used a probabilistic model to determine the karyotypic outcomes of multipolar divisions (see Materials and Methods and **Figure S2a**). We found that daughter nuclei emerging from multipolar divisions in tetraploid cells were very likely to bear a monosomy or nullisomy for at least one chromosome (**Figure 3a, S2b**). Nullisomic cells (and perhaps cells with certain monosomies) are expected to be unable to proliferate further. This was confirmed experimentally in long-term time lapse microscopy experiments (**Figure 3b**), which showed that cells produced by multipolar divisions were more likely to die or arrest compared to cells produced by bipolar divisions (52% vs. 10%; **Figure 3c**). Previous studies have shown that diploid and near-triploid cells undergoing multipolar divisions produce daughter cells that die^31^. Our data indicate that this may also be the case for tetraploid cells undergoing multipolar divisions.

**Figure 3.**
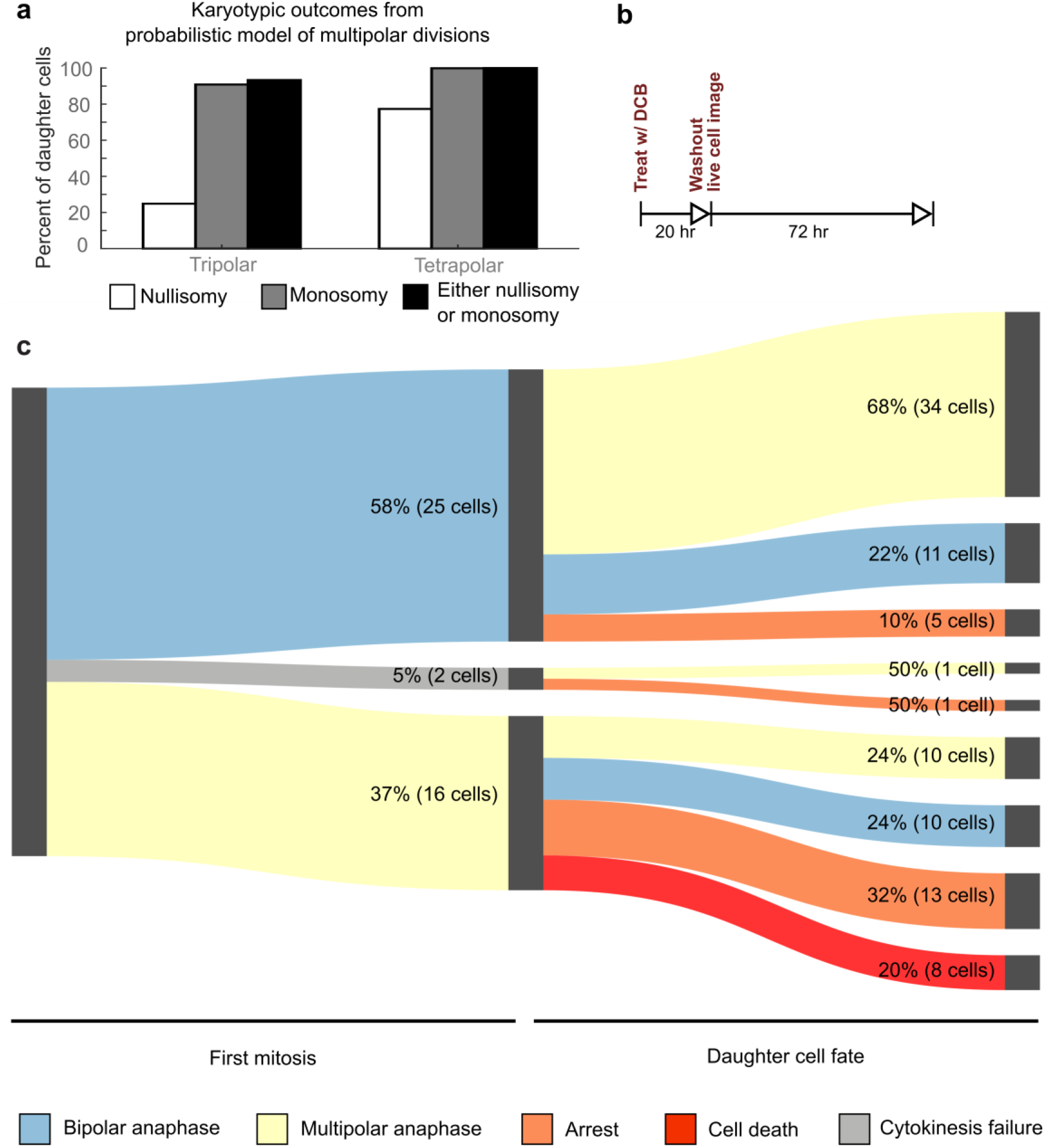
Daughters of multipolar divisions are likely to bear nullisomies or monosomies, explaining their likelihood to die or arrest over the subsequent 48 hours. (a) Results from probabilistic modeling to determine the likelihood of nullisomy and/or monosomy as a result of multipolar division. (b) Experimental design for live cell imaging to analyze the first two mitotic divisions of newly formed tetraploid cells. (c) Sankey diagram showing results of the 72 hr live cell imaging experiment in which newly formed tetraploid cells were followed through two consecutive cell divisions.

### Supernumerary centrosomes that arise through tetraploidization quickly disappear from the population

The observation that highly aneuploid cells make up only a very small fraction of the population by 12 days after tetraploidization (**Figure 2c-d**) suggests that the rate of multipolar divisions (which generate these cells) decreases over time. This could be due to either an increased ability of the extra centrosomes to cluster in the tetraploid cells or to elimination of the extra centrosomes. To explore this, we investigated if and how centrosome number varies over the same 12-day evolution period by analyzing cells immunostained for centrin immediately following cytokinesis failure and every two days thereafter (**Figure 4a**).

**Figure 4.**
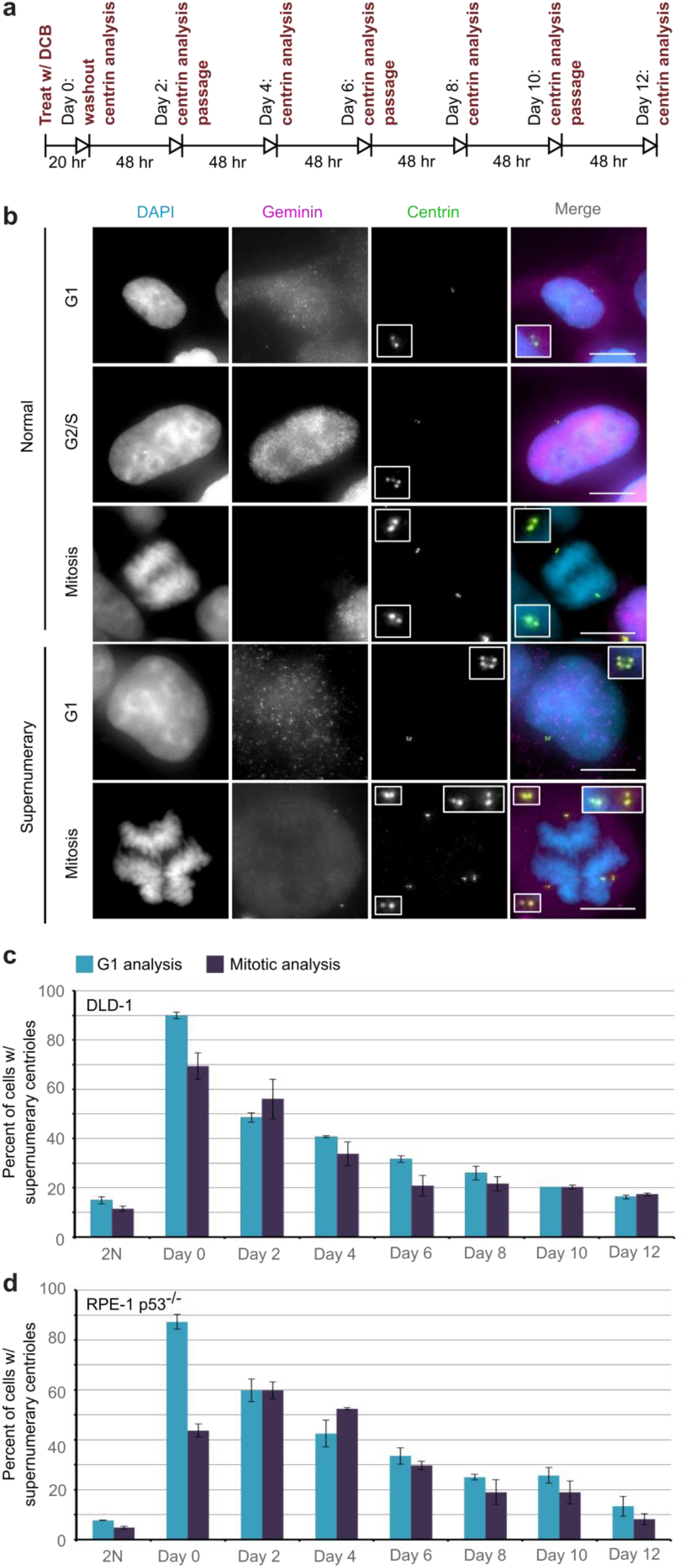
Extra centrosomes are rapidly lost from cell populations after experimental induction of tetraploidization by cytokinesis failure. (a) Experimental design for time course experiments designed to analyze centrosome number in newly formed tetraploid cells evolving over a 12-day period. (b) Examples of interphase and mitotic cells with normal centrosome number (top) or supernumerary (bottom) centrosomes. Scale bar, 10 μm. (c-d) 12-day time course analysis of centrosome number in newly formed tetraploid DLD-1 (c) and newly formed tetraploid RPE-1 p53^−/−^ (d) cells analyzed in both mitosis and G1 (cells negative for nuclear geminin staining). Centrosome number data are reported as mean ± S.E.M. from two biological replicates.

We performed this analysis in both mitotic and G1 cells and obtained similar results at all time points, except immediately following DCB washout (**‘Day 0’ in Figure 4c-d**). This discrepancy at day 0 could be explained by a delay in mitotic entry of 4N cells, particularly for the RPE-1 p53^−/−^ cells. Despite this difference at day 0, the trend was very clear. The fraction of the G1 cell population containing supernumerary centrioles after a 20 hr cytokinesis block was 90% and 87.3% in DLD-1 and RPE-1 p53^−/−^ cells, respectively. However, this fraction rapidly diminished over the 12-day observation period, reaching frequencies (16.3% and 13.3%, **‘Day 12’ in Figure 4c-d**) that are very close to the frequencies of cells with supernumerary centrioles in the parental populations (15% and 7.7%, respectively, in DLD-1 and RPE-1 p53^−/−^). Moreover, the number of cells with supernumerary centrioles at day 12 was substantially smaller than the number of cells with ~4N chromosome number (63% and 33%, respectively) at the same time point (**Figure 2**), indicating that a large fraction of the tetraploid cells that are present 12 days post cytokinesis failure have lost their extra centrosomes.

### Tetraploid cells can inherit a normal centrosome number through asymmetric centrosome clustering during cell division

We reasoned that one way in which tetraploid cells could regain a normal number of centrosomes while maintaining tetraploid chromosome numbers would be by asymmetrically clustering the centrosomes during formation of a bipolar spindle. As a result, one daughter cell would receive three centrosomes and the other daughter would receive one centrosome, but both would receive a chromosome number corresponding to ~4N.

To explore this possibility, we analyzed bipolar mitotic DLD-1 and RPE-1 p53^−/−^ cells fixed and immunostained for centrin immediately following washout of DCB (**Figure 5a-b**). We found nearly even numbers of bipolar DLD-1 cells with symmetric vs. asymmetric centrosome clustering in late mitosis (metaphase, anaphase, or telophase), while bipolar RPE-1 p53^−/−^ showed a moderate bias towards symmetric clustering of centrosomes (**Figure 5c**).

**Figure 5.**
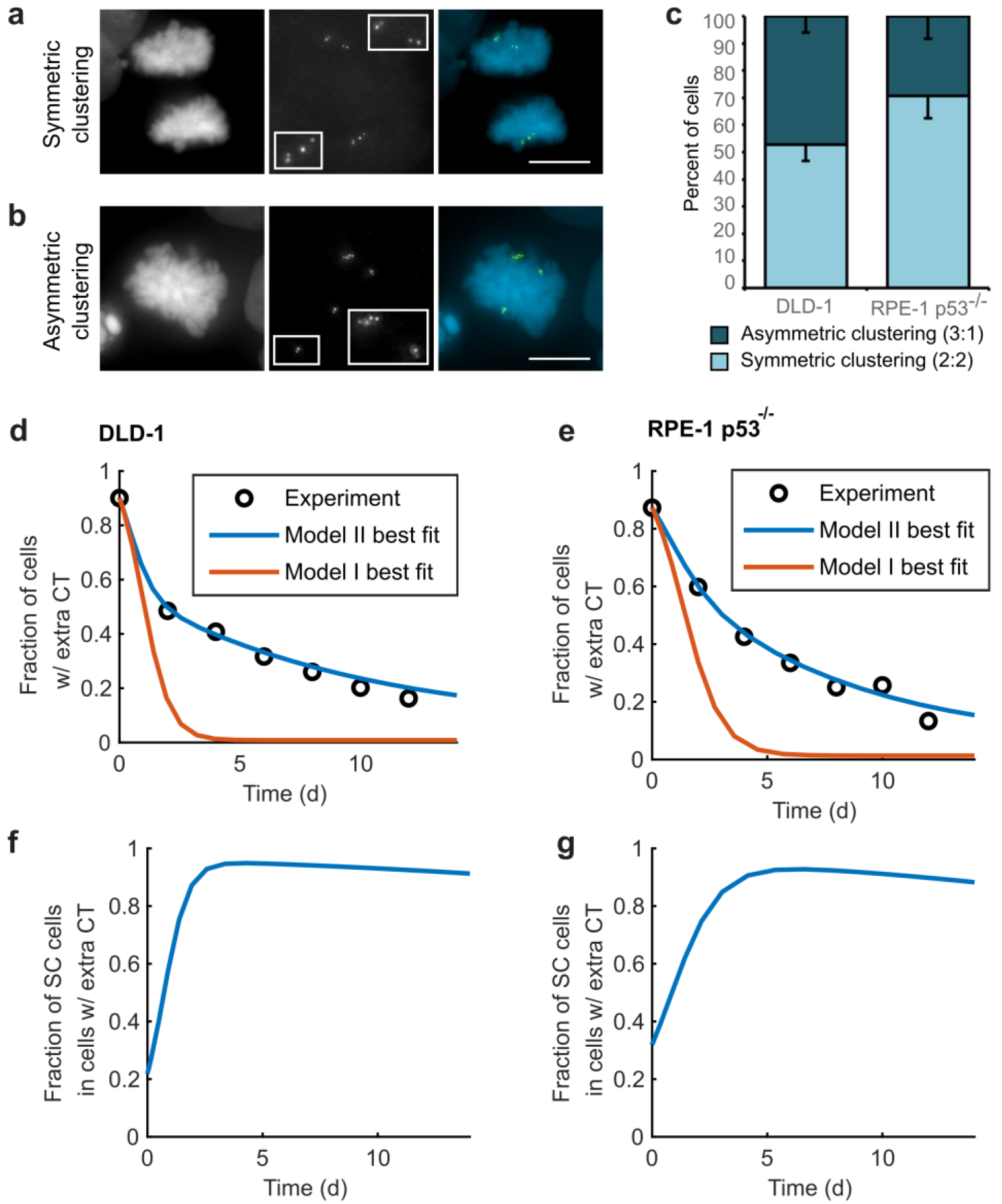
Asymmetric clustering of centrosomes in bipolar divisions can explain the formation of tetraploid cells with a normal centrosome number. (a) Example of anaphase cell with supernumerary centrosomes clustered symmetrically into a bipolar spindle, so that two centrosomes (four centrioles) are at each spindle pole. (b) Example of a late prometaphase cell with supernumerary centrosomes clustered asymmetrically into a near-bipolar spindle, so that one centrosome (two centrioles) is at one spindle pole and three centrosomes (six centrioles) are at the other spindle pole. Scale bar, 10 μm. (c) Quantification of symmetric vs. asymmetric centrosome clustering in newly formed tetraploid DLD-1 and RPE-1 p53^−/−^ cells. Data are reported as mean – S.E.M. from three biological replicates. (d-e) Modeling results for centrosome evolution based on Model I (red) or Model II (blue), superimposed over experimental data (circles), for DLD-1 (d) and RPE-1 p53^−/−^ (e) cells. When available, parameters that reflect experimental data were used (see supplementary modeling methods for further details and Table S1 for parameter values). (f-g) Fraction of cells with supernumerary centrosomes that are, over time, SC cells based on model II for DLD-1 (f) and RPE-1 p53^−/−^ (g) cells.

For asymmetric centrosome clustering to explain evolution of a tetraploid cell population with normal centrosome number, one would also have to assume that the daughter cell inheriting a single centrosome has a selective advantage over the daughter cell inheriting extra centrosomes (e.g., due to the likelihood of multipolar division in cells with extra centrosomes). To test this, we built a mathematical model based on this assumption (for model details, see Materials and Methods, **Table S1**, and **Figures S3-S6**). We started with a simple model in which, initially, 87-90% of the cells have two centrosomes in G1 (four in S/G2/M), corresponding to the experimentally observed frequencies (**Figure 4c-d**). Based on probabilities observed in experimental populations (**Figure 1b**), cells in the model have a different probability of dividing in a bipolar vs. multipolar fashion. The daughter cells from multipolar divisions were assumed to have reduced viability in line with our experimental observations (**Figure 3c**). The cells resulting from bipolar divisions were assumed to display different fates, depending on the number of centrosomes inherited. Cells inheriting two centrosomes would display the same fate as the initial cells; cells inheriting three centrosomes were assumed to undergo multipolar division with a much higher frequency; and cells inheriting a single centrosome were assumed to become stable cells that undergo bipolar divisions and display a low death rate. Although this initial model captured centrosome loss, it predicted centrosome loss over a much shorter time scale than was observed experimentally for DLD-1 and RPE-1 p53^−/−^ cells (**Figure 5d-e, red line**). The final fraction of cells with extra centrosomes predicted from the model was also substantially lower than what was experimentally observed. Parameter optimization within a reasonable range could not solve this discrepancy (**Figure S3b, S6a**). In particular, the final steady-state fraction of cells with extra centrosomes is strongly constrained by the probability of cells with normal centrosome number to fail cytokinesis (generating new cells with extra centrosomes, which counteracts supernumerary centrosome loss in the population) (**Figure S6a**). For the experimentally quantified^32^ probability (~2.5%) of spontaneous cytokinesis failure in DLD-1 cells, the steady-state fraction of cells with extra centrosomes cannot reach the observed value.

Closer examination of the experimental data (**Figure 3c**) indicated that there may be a sub-fraction of newly formed DLD-1 tetraploid cells that can cluster their centrosomes more efficiently than other cells, as indicated by the presence of cells that undergo two consecutive bipolar divisions right after tetraploidization (**Figure 3c**, two consecutive blue sections). These “super clustering” (SC) cells would tend to persist in the population and therefore predominantly make up the final, steady state population of tetraploid cells with supernumerary centrosomes (**Figure 5f-g**). When SC cells are included in the model, the model output was no longer constrained by the probability of cytokinesis failure (**Figure S6b**) and the final fraction of cells with extra centrosomes could match the experimentally observed values (**Figure 5d-e, blue line**).

Altogether, our modeling results show that asymmetric centrosome clustering, along with a selective advantage of cells that inherit a single centrosome, can be sufficient to explain the loss of extra centrosomes in newly formed tetraploid cells, leading to the evolution of cell populations with tetraploid chromosome numbers but normal centrosome numbers (i.e., 1 centrosome, 2 centrioles in G1 tetraploid cells).

### Long-term live-cell imaging confirms that centrosome elimination and stable tetraploid cells arise via asymmetric centrosome clustering and natural selection

To obtain definitive experimental evidence of the mechanism by which tetraploid cell populations lose their extra centrosomes, we generated cell lines stably expressing GFP-tagged centrin from both DLD-1 and RPE-1 p53^−/−^ cells and performed live cell imaging experiments. First, we quantified centrin dots in binucleate cells synchronized in prophase/early prometaphase by RO3306 washout (**Figure 6a**; see materials and methods for details). We found that all of the binucleate cells that we observed (n = 36) contained supernumerary centrosomes. Moreover, observation of the first mitotic division after tetraploidization showed that centrosomes were duplicated prior to mitosis and that they were never lost/extruded during mitosis (**Figure 6b**). These observations indicate that the rapid loss of supernumerary centrosomes evident in our time-course experiment (**Figure 4**) cannot be explained by suppression of centrosomes duplication, extrusion of centrosomes, or centrosome degradation.

**Figure 6.**
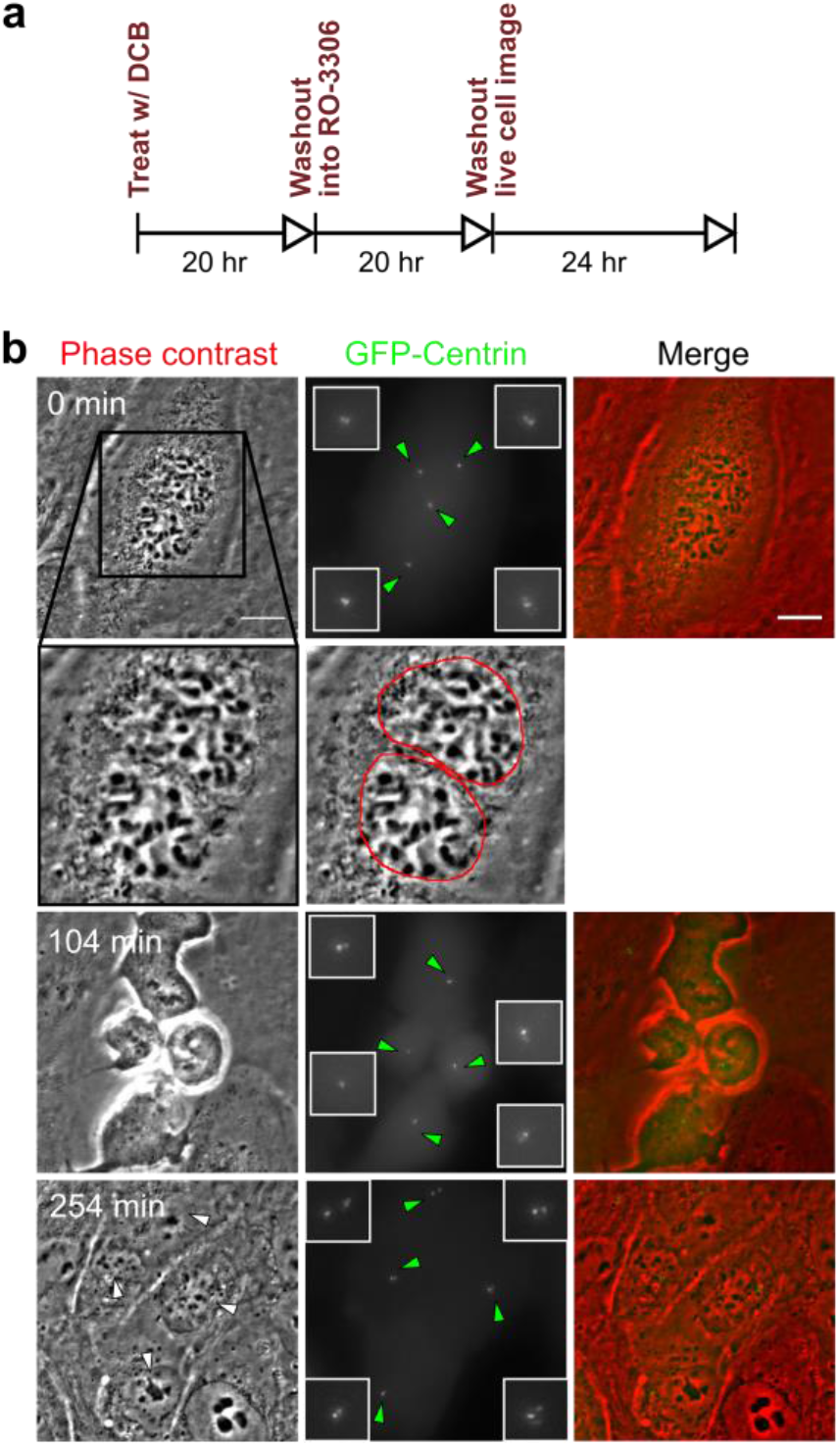
Extra centrosomes are not lost during cell division. (a) Experimental design for short term live cell imaging of binucleate cells expressing GFP-labeled centrin. Cells were tracked through their first cell division after induced cytokinesis failure. (b) Neither mitotic cells nor their daughters were found to have, in total, fewer centrosomes than their mother cell. Scale bar, 10 μm.

Next, we used these GFP-centrin cell lines to investigate the fates of daughter cells generated in the first tetraploid cell division in relation to the number of centrosomes they inherit. Because previous observations^20^ and our own data (**Figure 3c**) indicated that the progeny of multipolar divisions display reduced viability, and since only bipolar or near-bipolar divisions are likely to generate the evolved (12 days) near-tetraploid cell population observed in our time-course experiment, we focused on fates of daughter cells generated from bipolar divisions. We imaged newly generated tetraploid (binucleate) cells by phase contrast microscopy for 24 hrs, after which we determined the number of GFP-centrin dots present in the daughter cells. These cells were then imaged for an additional 48 hrs by phase contrast microscopy (**Figure 7a**). Consistent with our previous results (**Figure 5**), cells that divided in a bipolar manner showed a mix of symmetric and asymmetric centrosome clustering, without a strong preference for one mode (**Figure 7b-c**). For both DLD-1 and RPE-1 p53^−/−^ cells, the majority of cells that inherited a normal centrosome number (1 centrosome/ 2 centrioles) went on to divide in a bipolar manner. In contrast, cells that inherited too many centrosomes went through a mix of fates, dominated by multipolar divisions, arrest, and cell death. These data, together with our mathematical modeling, strongly suggest that populations of stably dividing tetraploid cells containing a normal number of centrosomes can arise through the genesis of cells with a normal number of centrosomes via asymmetric clustering of centrosomes (3:1) into bipolar spindles and selective pressure against cells with extra centrosomes.

**Figure 7.**
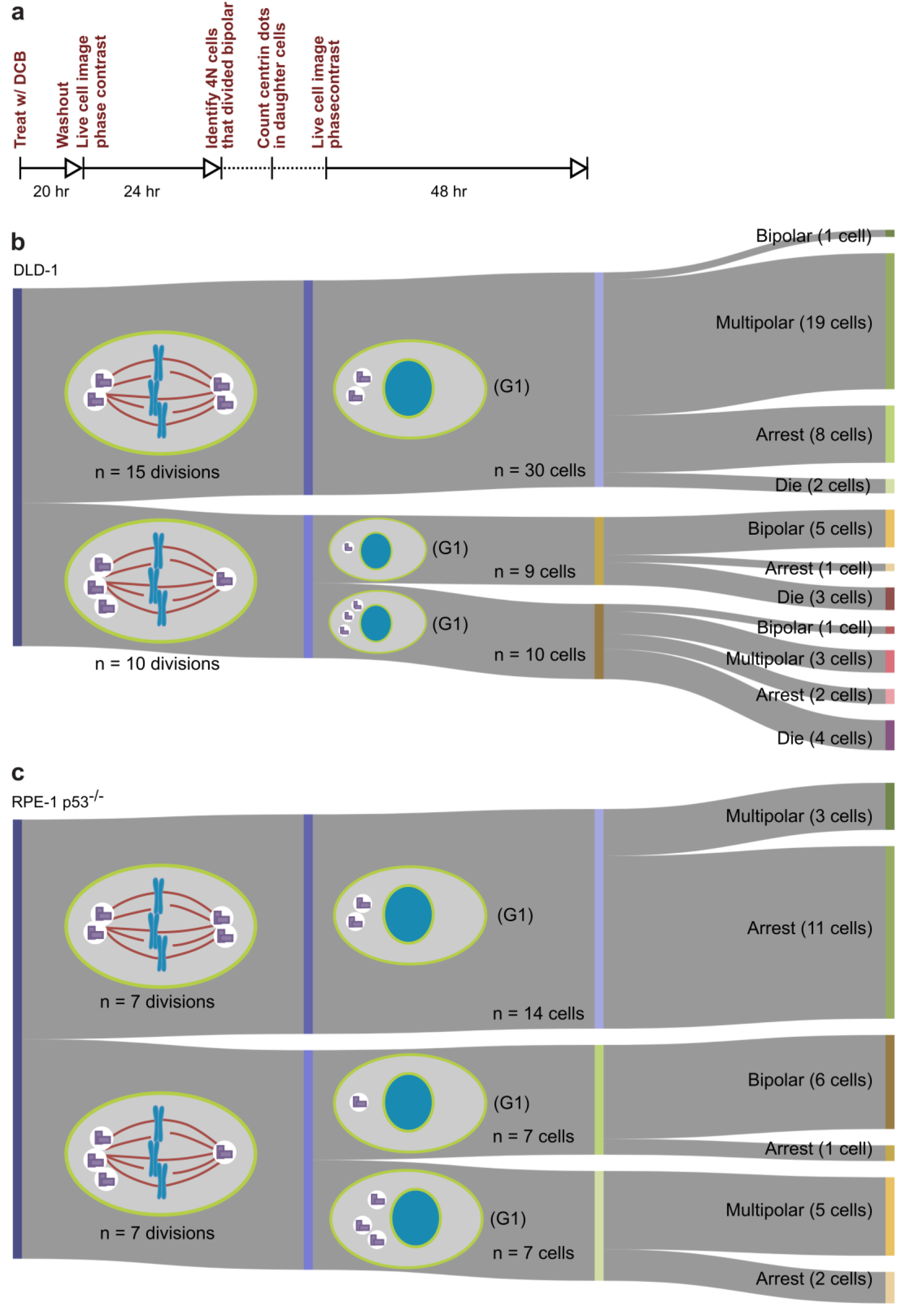
Cells that inherit a single centrosome are the only ones that continue to consistently divide in a bipolar manner. (a) Experimental design for long-term live cell imaging of cells with GFP-labeled centrin. (b) Data from long-term live cell imaging of newly formed DLD-1 tetraploid cells, expressing GFP-labeled centrin 2, in which the numbers of centrosomes inherited by daughter cells of bipolar divisions was tracked. (c) Data from long-term live cell imaging of newly formed RPE-1 p53^−/−^ tetraploid cells, expressing GFP-labeled centrin 2, in which the numbers of centrosomes inherited by daughter cells of bipolar divisions was tracked.

## Discussion

### Newly formed tetraploid cells rapidly lose the extra centrosomes while maintaining the extra chromosomes

Here, we show that populations of newly formed tetraploid cells rapidly evolve *in vitro* to retain a near-tetraploid chromosome number while losing the extra centrosomes gained at the time of tetraploidization. By combining fixed cell analysis, live cell imaging, and mathematical modeling, we show that this happens by a process of natural selection (**Figure 8**). Specifically, tetraploid cells that inherit a single centrosome during a bipolar division with asymmetric centrosome clustering are favored for long-term survival. Conversely, the majority of cells with extra centrosomes are eventually eliminated because of their high probability of undergoing a multipolar division, which would produce daughters with insufficient amounts of genetic material (**Figure 8**).

**Figure 8.**
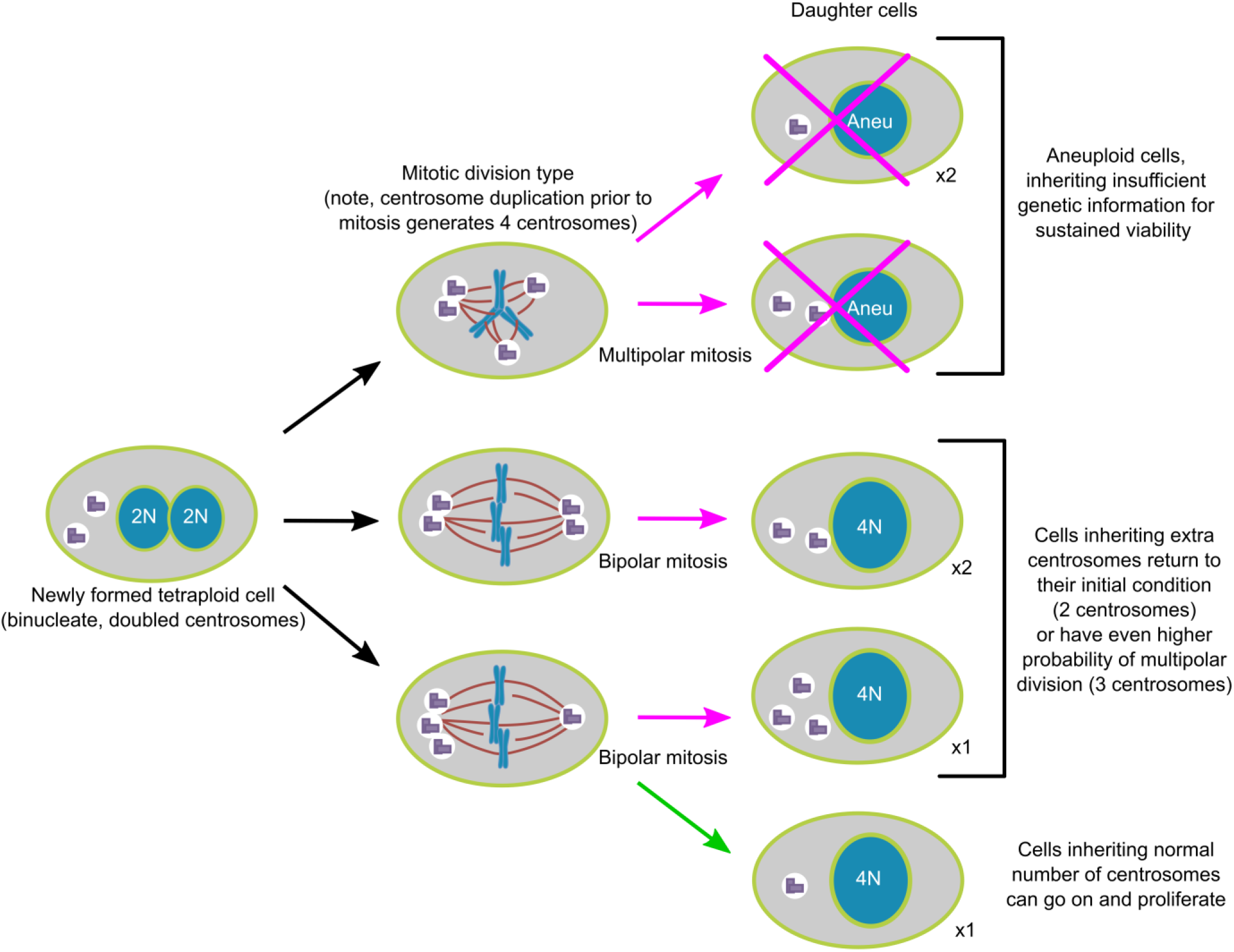
Asymmetric clustering of centrosomes defines the early evolution of tetraploid cells. The diagram illustrates the possible fates of a newly formed tetraploid cell and the mechanism by which tetraploid cells containing a normal number of centrosomes arise.

Our findings can explain previous anecdotal reports^20,26,27^ that clones isolated after experimental inhibition of cytokinesis consisted of tetraploid cells with a “normal” number (i.e., same number as in diploid cells) of centrosomes. Our study suggests that this pattern of centrosome number evolution after tetraploidization may be a general rule, rather than the exception. Our work also reveals the mechanism (**Figure 8**) by which tetraploid cells containing a normal number of centrosomes arise. Finally, our mathematical model frames tetraploid cell evolution (**Figure 6**) and may be used to better understand how tetraploidy contributes to tumor initiation and progression *in situ*.

### Tetraploidization and tumorigenesis: extra chromosomes vs. extra centrosomes as the causal link

The link between tetraploidization and tumorigenesis is supported by strong experimental evidence. Indeed, cancer genome sequencing data indicated that tetraploidization occurs at some point during the progression of a large fraction of tumors^13^. Moreover, tetraploid mouse epithelial cells were shown to be more tumorigenic than their non-tetraploid counterparts when injected in nude mice^14,15,33^. A popular model for how tetraploidy may promote tumorigenesis is that the extra centrosomes (which arise concomitantly with tetraploidization) contribute to cancer phenotypes^25^. This idea is supported by the following important observations: centrosome amplification is frequently observed in the pre-malignant stages of certain cancers^34^ and is observed in a large fraction of human tumors^35,36,37,38,39^, in which it correlates with poor prognosis/advanced disease stage^34,40^; extra centrosomes can promote tumorigenesis in mouse^23,24^ and enhance the invasive behavior of mammary epithelial cells grown in 3D cultures^22^; finally, supernumerary centrosomes promote chromosome mis-attachment and mis-segregation^20,21^, leading to chromosomal instability, a hallmark of cancer believed to drive tumor evolution^41^. Together, these observations indicate that extra centrosomes are likely to contribute to tumor initiation and/or progression.

Nevertheless, a number of studies and observations suggest that tetraploidy *per se* may promote the emergence of cancer phenotypes. For instance, tetraploidy was shown to increase tolerance for genomic changes, leading to the rapid evolution of complex genomes^42^, as seen in cancer and many tetraploid cells show increased chromosomal instability compared to diploid cells, even when no extra centrosomes are present^26^. Additionally, polyploid cells were shown to be more resistant than their diploid counterparts to oxidative stress, genotoxic insult, irradiation, and certain chemotherapeutic drugs^26,43,44^. Lastly, in cancer patients, genome-doubling in early stage tumors was shown to correlate with poor relapse-free survival^42^, although centrosome number was not examined in these patients.

In light of our findings, one could imagine that in certain instances, cells experiencing a genome doubling event may initially lose their extra centrosomes and then re-acquire them at a later time, depending on additional factors. At least one example in the literature provides evidence for such a series of events. In Barrett’s esophagus, a pre-malignant condition that predisposes to esophageal cancer^45,46,47^, a characteristic series of events has been described: loss of function of the p53 tumor suppressor gene occurs early in disease progression, followed by an accumulation of 4N cells prior to metaplasia, and finally, the accumulation of aneuploid cells as the tissue transitioned to metaplasia^10^. A different study reported centrosome amplification prior to the transition to metaplasia^48^, corresponding to the time when tetraploid cells accumulate^10^. Importantly, however, the frequency of supernumerary centrosomes decreased with transition to metaplasia and with subsequent neoplastic progression^48^. These results closely mirror the dynamics of evolution seen in our study and illustrate that extra centrosomes can be present early in tumor development (around the time when tetraploidy appears) but subsequently be lost. Therefore, while tetraploidy and supernumerary centrosomes are both individually linked with tumorigenesis, the link between tetraploidy, extra centrosomes, and disease progression may be less direct than conventionally thought.

Tetraploidization is intimately linked with the birth of extra centrosomes; however, tetraploidization may not lead to stable acquisition of supernumerary centrosomes unless (i) specific cellular/genetic changes have occurred (e.g., enhanced centrosome clustering^49,50^) to allow the cell to maintain its extra centrosomes and/or (ii) certain conditions in the tissue microenvironment exist that favor or necessitate the presence of extra centrosomes. Alternatively, tetraploid cells may initially lose their extra centrosomes and then re-acquire them at a later time, as a result of genome instability, which may lead to the non-stoichiometric production of proteins involved in centrosome duplication. The evolutionary pattern that newly formed tetraploid cells will follow may vary depending on many factors, including genetic background, functional requirements in a given tissue/organ, or a variety of extracellular factors (both physical and physiological).

## Acknowledgements

We would like to thank the labs of Neil Ganem and Tim Stearns for reagents and Meng-Fu Bryan Tsou for the RPE-1 p53^−/−^ cell line. Funding for this work was provided by the Virginia Tech Fralin Life Science Institute, ICTAS Center for Engineered Health, and College of Science (through a Dean’s Discovery fund award to D.C.). Work in the Cimini lab is further supported by NSF grant MCB-1517506 to D.C. We further acknowledge and thank the MultiSTEPS IGERT and BIOTRANS IGEP graduate programs for providing training and funding for N.C.B. and members of the Cimini and Hauf labs for helpful discussion.

