## Supplemental Material for "Asymmetric clustering of centrosomes defines the early evolution of tetraploid cells"

### Methods

#### *Experimental Methods*

##### Cell lines and culture conditions

DLD-1 cells (ATCC CCL-221) were purchased from the American Type Culture Collection (ATCC, Manassas, VA). The hTERT immortalized RPE-1 $p53^{-/-}$ cell line[S1] (referenced throughout the manuscript as RPE-1 $p53^{-/-}$) was a gift from Dr. Meng-Fu Bryan Tsou (Memorial Sloan Kettering Cancer Center). DLD-1 cells were cultured according to ATCC recommendations in RPMI 1640 media with ATCC modification (Thermo Fisher Scientific – Gibco, CA, USA) supplemented with 10% fetal bovine serum (FBS; Thermo Fisher Scientific – Gibco, CA, USA) and 1% antibiotic-antimycotic (Thermo Fisher Scientific – Gibco, CA, USA). RPE-1 $p53^{-/-}$ cells were cultured according to the ATCC recommendations for hTERT-immortalized RPE-1 cells in 1:1 mixture of DMEM/F-12 with HEPES (Thermo Fisher Scientific – Gibco, CA, USA) also supplemented with 10% FBS and 1% antibiotic-antimycotic. All cells were grown on tissue culture polystyrene flasks (Corning, Tewksbury, MA) and were maintained in a humidified incubator at 37°C and 5% $CO_2$.

Tetraploid DLD-1 and RPE-1 $p53^{-/-}$ cells were generated by treating diploid cell cultures with 1.5 µg/mL dihydrocytochalasin B (DCB; Sigma Aldrich, Saint Louis, MO) for 20 hrs. For live cell experiments, cells were washed out (4 times with 1X PBS) into imaging media and immediately taken to the microscope for imaging following.

##### Generating virally transduced cell lines

The GFP-Centrin 2 gene was PCR amplified from a modified pLL3.7 plasmid with unknown selection (gift of Tim Stearns, Stanford University), using forward and reverse primers designed to match the two termini of the fusion protein. The forward and reverse primers used (including restriction sites for NOTI and SALI and terminal non-sense nucleotides) were (with start and stop codons underlined):

(F) CAATAAAGCGGCCGCATGGTGAGCAAGGGCGAGGAGCTGT and

(R) GGACTGGTGGTCTGCGTCGACTTAATAGAGGCTGGTCTTTTTCATG.

Cleaned PCR product was ligated into the pLNXC2 retroviral expression vector by directional cloning using NOTI and SALI restriction enzymes. The presence of GFP-centrin 2 gene in plasmid DNA was confirmed by restriction digests visualized on DNA gels and via transient transfection into RPE-1 $p53^{-/-}$ cells to confirm centrosomal localization. GFP-Centrin expressing DLD-1 and RPE-1 $p53^{-/-}$ cells were generated by transduction with retroviral particles. GP-293 cells containing retroviral *gag* and *pol* genes (ClonTech Laboratories Inc., Mountain View, CA #631458) were co-transfected with the expression vector and the pVSV-G plasmid (Addgene, Cambridge, MA). 48 hrs after transfection, supernatant was collected, filtered through a 0.45 µm pore (GD/X sterile 0.45 µm CA filter, GE Whatman PLC, Pittsburgh, PA), mixed with polybrene (Sigma-Aldrich, Saint Louis, MO) at a final concentration of 10 µg/ml, and added directly to the cells. After 24 hrs, cell media was replaced with fresh culture media. Starting 72 hrs after viral transduction, transduced cells were selected with with G418 at a concentration of 500 µg/ml until negative control cells (untransduced cells treated with the same concentration of antibiotic) were dead, or approximately two weeks.

Cells co-expressing RFP-H2B were generated by further transducing GFP-Centrin 2 expressing cells, via the protocol described previously, using a pBABE retroviral plasmid containing RFP-H2B and a puromycin selection gene (gift from Neil Ganem, Boston University). Transduced cells were selected with puromycin at a concentration of 5 µg/ml (RPE-1 $p53^{-/-}$) or 3.8 µg/ml (DLD-1).

**Phase contrast live cell microscopy**

For live-cell experiments, all cells were grown on MatTek glass bottom dishes with No. 1.5 glass (MatTek Corporation, Ashland, MA). At the time of imaging, cell media was replaced with L-15 media supplemented with 4.5 g/l glucose (high glucose). All live cell experiments were performed on a Nikon Eclipse Ti inverted microscope (Nikon instruments Inc., NY, USA) equipped with phase-contrast trans-illumination, transmitted light shutter, ProScan automated stage (Prior Scientific, Cambridge, UK), CoolSNAP HQ2 CCD camera (Photometrics, AZ, USA), Lumen200PRO light source (Prior Scientific, Cambridge, UK), and a temperature and humidity controlled incubator (Tokai Hit, Japan). For 24 hr and 72 hr live cell phase contrast videos, images were acquired every 6 minutes through a 20X/0.3 NA A Plan corrected phase contrast objective for the duration of the experiment. Time-lapse videos were analyzed using NIS Elements AR software (Nikon Instruments Inc., NY, USA) to determine the nature of division (bipolar, tripolar, tetrapolar) at anaphase and the subsequent number of daughter cells formed after cytokinesis.

**Time course experimental procedure**

Time course (12 day) experiments were performed by seeding all cells needed for the first two time points (day 0 and day 2) along with a flask designated for propagating the experiment. For each replicate for DLD-1 cells, this included T-25 flasks seeded with $1x10^6$ (day 0 metaphase spreads) and $5x10^5$ (day 2 metaphase spreads), a T-75 flask with $1x10^6$ cells, and acid-washed coverslips inside 35 mm Petri dishes with $2.5x10^5$ (day 0) and $1x10^5$ (day 2) cells for combined centrin/ geminin immunostaining. On day 2, the T-75 flask was used to seed cells for the next two time points as follows: two T-25 flasks (metaphase spreads), one T-75 flask (propagating), and coverslips (centrin/ geminin immunostaining). This was repeated for the entire 12-day period. The experiment was designed in the same way for RPE-1 $p53^{-/-}$ cells, but cell counts were as follows: T-25 flasks seeded at $1x10^6$ cells (earlier time point, e.g. day 0) and $5x10^5$ (later time point, e.g. day 2); T-75 seeded at $1.5x10^6$ cells; coverslips seeded at $1.25x10^5$ (earlier time point) and $8.5x10^4$ cells (later time point).

**Chromosome spread preparation and analysis**

Cell cultures were grown in T-25 flasks as described in the previous section. For chromosome spread preparation, cells were incubated in their respective media containing 50 ng/ml colcemid (Invitrogen – Karyomax, Waltham, MA) at 37°C for 5 hr to enrich for mitotically arrested cells. The cells were then collected by trypsinization and centrifuged at 1000 rpm for 5 min. Pre-warmed (37°C) hypotonic solution (0.075 M KCl) was added drop-wise to the cell pellet and incubated for 18 (DLD-1 cells) or 15 (RPE-1 $p53^{-/-}$ cells) minutes at 37°C. Several drops of freshly prepared fixative (3:1 methanol:glacial acetic acid) were added before centrifugation at 1000 rpm for 5 min. Supernatant was aspirated, fixative was added dropwise, and the cell suspension was again centrifuged at 1000 rpm for 5 min. The fixation step was repeated two more times and fixed cells were finally dropped on microscope slides. For RPE-1 p53$^{-/-}$ cells, a homemade humidity chamber constructed from PVC piping, plastic sheeting, and a nebulizer was used when drying slides to improve chromosome spread quality (effect of humidity on chromosome spread quality was described previously[S2]). Chromosome spreads were then stained with 300 nM DAPI (Thermo Fisher Scientific – Invitrogen, Waltham, MA) for 10 minutes. DAPI-stained slides were mounted with an antifade solution containing 90% glycerol and 0.5% N-propyl gallate and sealed under a 22x50 mm coverslip (Corning Incorporated, Corning, NY) with nail polish. For chromosome counting, images of DAPI-stained chromosome spreads were acquired with the Nikon Eclipse Ti inverted microscope setup described in an earlier section. Images were acquired using either a 60X/1.4 NA or a 100X/1.4 NA Plan-Apochromatic phase contrast objective. After image acquisition, chromosome spreads were visualized in gray scale and chromosomes were counted using the count function in NIS elements.

**Cell death assays**

To measure cell death, $5x10^4$ cells were plated in each of three wells of a 6-well plate and $1x10^6$ cells were plated in a T-25 flask. The following day, cells were treated with 1.5 µg/ml DCB for 20 hr. After 20 hr, day 0 cells' supernatant was collected, while the adherent cells were washed (3 times using PBS) and harvested in trypsin. The supernatant, all the washes, and the cell suspension were collected in the same tube, centrifuged, and re-suspended in 400 µl PBS for counting. Viable cells were differentiated from dead cells by trypan blue exclusion. The numbers of living and dead cells were counted and the fraction of dead cells out of the total number of cells was calculated. Cell counting was performed on days 0, 1, 2, and every 2 days for the remainder of the 12-day period (with new wells being seeded from T-25 flasks on days 2, 6, and 10). Cell culture medium was changed 24 hrs before counting each day in order to only count cells that died within a 24 hr period.

**Immunofluorescence staining, image acquisition and data analysis**

For centrin and geminin immunostaining, cells were grown on sterilized acid-washed glass coverslips inside 35 mm Petri dishes. Cells were fixed in freshly prepared 4% paraformaldehyde in PHEM buffer (60 mM Pipes, 25 mM HEPES, 10 mM EGTA, 2 mM $MgSO_4$, pH 7.0) for 20 min at room temperature and then permeabilized for 10 min at room temperature in PHEM buffer containing 0.1% Triton-X 100. Following fixation and permeabilization, cells were washed three times with PBS and then blocked with 20% boiled goat serum (BGS) for 1 hr at room temperature. Cells were then incubated at 4°C overnight with primary antibodies diluted in 10% BGS. Next, cells were washed in PBS-T (PBS with 0.05% Tween 20) three times, and incubated at room temperature for 45 min with secondary antibodies diluted in 10% BGS. Cells were then washed four times with PBS-T, stained with DAPI (300 nM, Thermo Fisher Scientific – Invitrogen, Waltham, MA) for 5 min, and coverslips were mounted on microscope slides in an antifade solution containing 90% glycerol and 0.5% N-propyl gallate. Primary antibodies were diluted as follows: rabbit anti-geminin (Abcam, Cambridge, MA), 1:100; mouse anti-centrin (Abnova, Zhongli, Taiwan), 1:100. Secondary antibodies were diluted as follows: Rhodamine Red-X goat anti-rabbit (Jackson ImmunoResearch Laboratories, Inc., PA, USA), 1:100; Alexa 488 goat anti-mouse (Molecular Probes, Life Technologies, CA, USA), 1:200.

Centrin-stained samples were analyzed on a Nikon Eclipse TE2000 inverted microscope equipped with a 100X/1.4 NA Plan-Apochromatic phase contrast objective lens, motorized ProScan stage (Prior Scientific, Cambridge, UK), appropriate filter sets, and an XCITE 120Q light source (Excelitas

Technologies, Waltham, MA, USA). Analysis was performed visually in both interphase cells and mitotic cells. The number of centrin dots was counted in cells that were determined to be in mitosis by DAPI staining. Mitotic cells with four centrin dots (i.e., two dots corresponding to each centrosome of a bipolar spindle) were categorized as normal; mitotic cells with greater than four dots were categorized as possessing supernumerary centrosomes. Interphase analysis was performed in G1/G0 cells, as determined by absence of nuclear geminin staining[S3]. G0/G1 cells with two adjacent centrin dots (corresponding to a single centrosome) were classified as normal, whereas cells with greater than two centrin dots were classified as possessing supernumerary centrosomes. For centrosome clustering analysis, bipolar metaphase, anaphase, or telophase cells were analyzed for the number of centrin dots present at respective spindle poles. Representative z-stack image examples were acquired on the Nikon Eclipse Ti inverted microscope setup described in an earlier section. Images were acquired using either a 60X/1.4 NA or a 100X/1.4 NA Plan-Apochromatic phase contrast objective and appropriate filters.

For analysis of genome distribution in bipolar and multipolar divisions, images of ana-/telophase cells were acquired with a swept field confocal system (Prairie Technologies, WI, USA) on the same Nikon Eclipse TE2000-U inverted microscope described previously (Nikon Instruments Inc., NY, USA). The microscope was equipped with a CoolSNAP HQ2 CCD camera (Photometrics, AZ, USA), a multiband pass filter set (illumination at 405, 488, 561, and 640 nm), and an Agilent monolithic laser combiner (MLC400) controlled by a four channel acousto-optic tunable filter. Images were obtained by acquiring Z-stacks with 0.6 µm steps (Nyquist matched) so that the entire 3-D volume of the DNA was captured. Images were shading corrected using the NIS Elements shading correction function. Z-stacks were summed using the FIJI sum slices function. The freehand selection tool was used to trace the signal area corresponding to an ana-/telophase chromosome cluster and the percentage of the signal intensity relative to total DNA for an ana-/telophase cell was determined. To calculate the symmetry score, the ratio between the actual fluorescence intensity percentage and the expected signal intensity percentage for an even distribution to 2 (50%), 3 (33.3%) or 4 (25%) poles (depending on the polarity of the division) was first calculated for each chromosome cluster. Then, the standard deviation of all measurements for a cell was calculated as a 'symmetry score' (ss). If a division was perfectly symmetrical, ss = 0 and any ss > 0 indicates proportional increases in the asymmetry of DNA distribution to the poles.

**Live cell imaging of fluorescently labeled cells**

For live cell imaging of GFP-Centrin expressing cells, imaging was performed with a 60X/1.4 NA Plan-Apochromatic phase contrast objective lens (for RPE-1 $p53^{-/-}$ cells) or a 100X/1.4 NA Plan-Apochromatic phase contrast objective lens (for DLD-1 cells) controlled by Nikon Perfect Focus (Nikon Instruments Inc., NY, USA). In preparation for short-term live imaging of binucleate cells expressing GFP-centrin and RFP-H2B, the cells were washed out of DCB into media containing 9 µM of the CDK1 inhibitor RO3306 to arrest cells at the G2/M transition. After 4 hrs, the cells were again washed out into high glucose L-15 media lacking phenol red. Imaging was performed by identifying individual binucleate cells in prophase or early prometaphase using RFP-H2B signal. Cells were imaged at the home Z-position in phase contrast every 4 minutes and the FITC channel every 4 minutes with asymmetrical Z-stacks defined by the home position and a range of -2.4 µm and +5.8 µm in 0.6 µm steps (14 steps). Chromosomes were imaged by phase contrast instead of fluorescence (RFP) to keep illumination, and hence photodamage, to a minimum, given that phase contrast imaging required lower exposure times than fluorescence imaging. Cells were imaged for a total of 3-

4 hrs. The videos were then analyzed to determine the number of centrin dots (centrioles) in the early mitotic cells and again in the resulting daughter cells after division.

For long-term cell fate experiments (**Figure 7**), GFP-Centrin expressing cells were used. Binucleate cells were imaged at 10 minute intervals for 24 hrs via phase contrast microscopy under a 60X/1.4 NA or 100X/1.4 NA Plan-Apochromatic phase contrast objective lens. Following this period, a number of daughter cells were selected and the number of centrioles was quickly counted for each by eye. A phase contrast image was obtained, along with asymmetric Z-stack images in the FITC channel, defined by the home position and a range of -2.4 μm and +5.8 μm in 0.6 μm steps. These daughter cells were then tracked via phase contrast microscopy at 10-minute intervals for an additional 48 hr period to determine their subsequent fate.

*Modeling methods*

##### 1. Probabilistic model for karyotypic outcomes of multipolar divisions

Below we indicate how the probabilities of nullisomy and monosomy in tripolar or tetrapolar divisions were derived for a mother cell with $M$ unique chromosomes ($M$ = 23 for human cells), each with $k$ homologous copies before duplication (i.e., $k$-ploid). For simplicity, we made the following assumptions:

1. The possibility of chromosome missegregation is ignored. Sister chromatids from each chromosome are partitioned to different spindle poles and end up in different daughter cells. The chromosome partitioning is otherwise random.
2. Homologous chromosomes are partitioned in the same way and independent of one another.

Due to the second assumption, the probability of an event (e.g., nullisomy, monosomy, or nullisomy/monosomy) in a daughter cell reads as Eq.(1). Because all sets of homologous chromosomes are equivalent in partitioning, the probability can be expressed in terms of the probability for Chr 1 without loss of generality.

$$\begin{aligned}\mathrm{P}\left(\text{event in the cell}\right) &= 1-\prod_{m=1}^{M}\left[1-\mathrm{P}\left(\text{event in the } m\text{-th chromosome in the cell}\right)\right] \\ &= 1-\left(1-\mathrm{P}\left(\text{event in Chr 1 in the cell}\right)\right)^{M}\end{aligned} \tag{1}$$

Next, we need to determine the probabilities of each event of interest for Chr 1, and use Eq.(1) to deduce the corresponding probabilities in the cell.

###### 1.1 Probability of nullisomy

Because sister chromatids have to be partitioned to different poles, the total number of equal ways to partition one pair of sister chromatids to $p$ poles reads as:

$$N_{1\times 2\to p} = \binom{p}{2} \tag{2}$$

where the bracketed expression represents the binomial coefficient.

Because sister chromatids from each chromosome are independent of each other in the partitioning, the total number of equal ways to partition $k$ pairs of sister chromatids to $p$ poles reads as:

$$N_{k\times 2\to p} = \binom{p}{2}^{k} \tag{3}$$

If any given pole receives 0 chromatids (i.e., nullisomy), then the total number of equal ways to partition $k$ pairs of sister chromatids to the remaining $p$-1 poles reads as:

$$N_{k\times 2\to p-1} = \binom{p-1}{2}^{k} \tag{4}$$

Hence, the probability that any given pole and the corresponding daughter cell bears a nullisomy for Chr 1 reads as:

$$\mathrm{P}\left(\text{nullisomy in Chr 1 in the cell}\right)=\frac{N_{k\times 2\to p-1}}{N_{k\times 2\to p}}=\frac{\left(\dfrac{(p-1)!}{(p-3)!2!}\right)^k}{\left(\dfrac{p!}{(p-2)!2!}\right)^k}=\left(\frac{p-2}{p}\right)^k \tag{5}$$

Note that the probability in Eq.(5) is not multiplied by another factor *p* for the number of possible poles/daughter cells, because we are looking for the probability of nullisomy of Chr 1 in a given daughter cell rather than in a given a cell division.

Plugging Eq.(5) into Eq.(1) yields the probability of nullisomy in a cell.

$$\mathrm{P}\left(\text{nullisomy in the cell}\right)=1-\left(1-\left(\frac{p-2}{p}\right)^k\right)^M \tag{6}$$

Plugging $M$ = 23, $k$ = 4, $p$ = 3 or 4 into Eq.(6) yields the results presented in **Figure 3a**. The number of nullisomies in a cell follows a binomial distribution $B(M,q)$, where $q=((p-2)/p)^k$ according to Eq.(5). The corresponding probability distribution for $M$ = 23, $k$ = 4, $p$ = 3 or 4 is plotted in **Figure S2b** (top).

##### 1.2 Probability of monosomy

If any given pole receives 1 chromatid (i.e., monosomy), then the total number of equal ways to partition the chromosomes reads as:

$$N=\underbrace{k}_{\substack{\text{choose 1 chr. pair}\\ \text{out of } k \text{ pairs}}}\times\underbrace{(p-1)}_{\substack{\text{choose 1 pole out of}\\ \text{the rest } (p-1) \text{ poles}\\ \text{for the chosen chr. pair}}}\times\underbrace{N_{(k-1)\times 2\to p-1}}_{\substack{\text{partition the rest } (k-1) \text{ chr. pairs}\\ \text{onto the rest } (p-1) \text{ poles}}}=k(p-1)\begin{pmatrix}p-1\\2\end{pmatrix}^{k-1} \tag{7}$$

Hence, the probability that any given pole and the corresponding daughter cell bears a monosomy for Chr 1 reads as:

$$\mathrm{P}\left(\text{monosomy in Chr 1 in the cell}\right)=\frac{N}{N_{k\times 2\to p}}=\frac{k(p-1)\left(\dfrac{(p-1)!}{(p-3)!2!}\right)^{k-1}}{\left(\dfrac{p!}{(p-2)!2!}\right)^k}=\frac{2k(p-2)^{k-1}}{p^k} \tag{8}$$

Plugging Eq.(8) into Eq.(1) yields the probability of monosomy in a cell.

$$\mathrm{P}\left(\text{monosomy in the cell}\right)=1-\left(1-\frac{2k(p-2)^{k-1}}{p^k}\right)^M \tag{9}$$

Plugging $M$ = 23, $k$ = 4, $p$ = 3 or 4 into Eq.(9) yields the results presented in **Figure 3a**. The number of monosomies in a cell follows a binomial distribution $B(M,q)$, where $q = 2k(p-2)^{k-1} / p^k$ according to Eq.(8). The corresponding probability distribution for $M$ = 23, $k$ = 4, $p$ = 3 or 4 is plotted in **Figure S2b** (bottom).

##### 1.3 Probability of nullisomy or monosomy

Because nullisomy and monosomy are mutually exclusive events, the probability that any given pole and the corresponding daughter cell bears either nullisomy or monosomy for Chr 1 reads as:

$$\begin{aligned} &\mathrm{P}(\text{nullisomy or monosomy in Chr 1 in the cell}) \\ &= \mathrm{P}(\text{nullisomy in Chr 1 in the cell}) + \mathrm{P}(\text{monosomy in Chr 1 in the cell}) \\ &= \frac{(p-2)^k + 2k(p-2)^{k-1}}{p^k} \end{aligned} \tag{10}$$

Plugging Eq.(10) into Eq.(1) yields the probability of nullisomy or monosomy in a cell.

$$\mathrm{P}(\text{nullisomy or monosomy in the cell}) = 1 - \left(1 - \frac{(p-2)^k + 2k(p-2)^{k-1}}{p^k}\right)^M \tag{11}$$

Plugging $M$ = 23, $k$ = 4, $p$ = 3 or 4 into Eq.(11) yields the results presented in **Figure 3a**.

#### 2. Model for centrosome number evolution in a cell population

##### 2.1 Model I

Model I was constructed based on the minimal assumptions about how centrosome numbers vary during cell divisions (**Figure S3a**). In other words,

1. A cell with normal centrosome number ($C_2$) undergoes normal division with probability $q$ and cytokinesis failure (→ $C_4$) with probability 1-$q$;
2. A cell with double centrosome number ($C_4$) undergoes bipolar division with probability $p$ and multipolar division with probability 1-$p$;
3. The bipolar division occurs in the symmetric fashion ($C_4$+$C_4$) with probability $r$ and in the asymmetric fashion ($C_2$+$C_6$) with probability 1-$r$; (note, the subscripts refer to the centrosome number of these daughter cells at the subsequent mitosis)
4. Multipolar division of an $C_4$ cell goes by 2$C_2$+$C_4$ with probability $s$ and 4$C_2$ with probability 1-$s$;
5. Multipolar division of an $C_4$ cell in the fashion of 4$C_2$ is fatal;
6. Multipolar division of a $C_4$ cell in the fashion of 2$C_2$+$C_4$ only has $C_4$ viable (equivalent to a normal $C_4$) with probability $f$.

In addition,

1. $C_2$ cells divide with rate $b_{C2}$, and die with rate $d_{C2}$;
2. $C_4$ cells divide with rate $b_{C4}$, and die with rate $d_{C4}$;
3. $C_6$ cells divide in multipolar fashion and die (there might be a small probability of viable division, which is neglected).

Based on the cell fate depicted in **Figure S3a**, the population dynamics are governed by the following ODEs:

$$\frac{dC_2}{dt} = b_{C2}(2q-1)C_2 + b_{C4}p(1-r)C_4 - d_{C2}C_2 \tag{12}$$

$$\frac{dC_4}{dt} = b_{C2}(1-q)C_2 + b_{C4}(2pr + (1-p)fs - 1)C_4 - d_{C4}C_4 \tag{13}$$

$$\frac{dC_6}{dt} = b_{C4}p(1-r)C_4 - d_{C6}C_6 \tag{14}$$

with initial condition $C_2(0) = \alpha N, C_4(0) = (1-\alpha)N, C_6(0) = 0$. The initial condition reflects the experimental observation that the newly induced 4N cell populations usually contain a certain fraction ($\alpha$) of $C_2$ (2N) cells.

Parameter sensitivity analysis (**Figure S3b**) indicated that the final fraction of cells with extra centrosomes strongly depends on $q$, the probability of cytokinesis failure in cells with normal centrosome number. In fact, the range of possible values for this final fraction is strongly constrained by the value of $q$ (see later discussion in section 2.2), even if choice of all parameters could span a wide range, instead of assuming fixed values as shown in Table S1. This is because the cytokinesis failure causes formation of new cells with extra centrosomes, and hence a large probability of cytokinesis failure maintains a higher fraction of these cells in the population.

When the cell division rate is sufficiently large compared to cell death rate in Eqs.(12)~(14), the number of cells in each type will increase infinitely (**Figure S4**, left column). This case does reflect the experiments, in which the cell cultures were regularly sampled and re-populated on fresh medium, effectively creating a finite sample of the unlimited population growth. Although the total population grows infinitely, the fractions of each cell type approach fixed steady state values (**Figure S4**, right column). In fact, the steady state fraction of each cell type can be analytically solved as shown in Section 2.3 below.

##### 2.2 Model II (with SC cells)

In the updated model (**Figure S5a**), we added SC cells, which are $C_4$ cells that can effectively cluster extra centrosomes, and regularly undergo bipolar division. For this new cell type, we made the following assumptions.

1. Cytokinesis failure in cells with normal centrosome number results in SC cells with probability, *v*.
2. SC cells divide symmetrically (SC+SC) with a probability, $r_S$. Otherwise, they divide asymmetrically ($C_2$+$C_6$).

3. SC cells have the same division and death rates as cells with normal centrosome number, because they are supposedly stable.

Based on the cell fate depicted in **Figure S5a**, the population dynamics are governed by the following ODEs:

$$\frac{dC_2}{dt} = b_{C2}\left(2q-1\right)C_2 + b_{C4}p\left(1-r\right)C_4 + b_{C2}\left(1-r_S\right)SC - d_{C2}C_2 \qquad (15)$$

$$\frac{dC_4}{dt} = b_{C2}\left(1-q\right)\left(1-v\right)C_2 + b_{C4}\left(2pr+\left(1-p\right)fs-1\right)C_4 - d_{C4}C_4 \qquad (16)$$

$$\frac{dSC}{dt} = b_{C2}\left(1-q\right)vC_2 + b_{C2}\left(2r_S-1\right)SC - d_{C2}SC \qquad (17)$$

$$\frac{dC_6}{dt} = b_{C4}p\left(1-r\right)C_4 + b_{C2}\left(1-r_S\right)SC - d_{C6}C_6 \qquad (18)$$

Parameter sensitivity analysis (**Figure S5b**) indicated that, based on this model, the final fraction of cells with extra centrosomes is most sensitive to $r_S$, the probability of symmetric division in SC cells, followed by $q$, the probability of cytokinesis failure in $C_2$ cells, and $v$, the probability of getting SC cells upon cytokinesis failure. While the initial model showed a strong constraint on $q$ (**Figure S6a**), the strength of this constraint is relaxed in the Model II (**Figure S6b**). The major constraint is now shifted to $r_S$ (**Figure S6c**), because asymmetric division (with probability 1- $r_S$) controls the conversion of SC cells back to $C_2$ cells. Nevertheless, ~90% probability of symmetric division is sufficient to maintain 20% cells with extra centrosomes in the steady state population.

##### 2.3 Steady state of cell fractions

Systems of homogenous linear ODE equations like Eqs.(12)~(14) and Eqs.(15)~(18) can be written in vector form as

$$\frac{d\mathbf{X}}{dt} = \mathbf{P}\cdot\mathbf{X} \qquad (19)$$

where $\mathbf{X} = \left(X_1, X_2, \ldots, X_N\right)$ is the list of variables.

The coefficient matrix, **P**, has the rate constants as entries. For Model I governed by Eqs.(12)~(14),

$$\mathbf{P} = \begin{bmatrix} b_{C2}\left(2q-1\right)-d_{C2} & b_{C4}p\left(1-r\right) & 0 \\ b_{C2}\left(1-q\right) & b_{C4}\left(2pr+\left(1-p\right)fs-1\right)-d_{C4} & 0 \\ 0 & b_{C4}p\left(1-r\right) & -d_{C6} \end{bmatrix} \qquad (20)$$

Likewise, for Model II governed by Eqs.(15)~(18),

$$\mathbf{P} = \begin{bmatrix} b_{C2}(2q-1)-d_{C2} & b_{C4}p(1-r) & b_{C2}(1-r_S) & 0 \\ b_{C2}(1-q)(1-v) & b_{C4}(2pr+(1-p)fs-1)-d_{C4} & 0 & 0 \\ b_{C2}(1-q)v & 0 & b_{C2}(2r_S-1)-d_{C2} & 0 \\ 0 & b_{C4}p(1-r) & b_{C2}(1-r_S) & -d_{C6} \end{bmatrix} \quad (21)$$

If $\det(\mathbf{P}) \neq 0$, then Eq.(19) only has the trivial steady state where all variables equal zero. This trivial steady state is unstable if the overall proliferation rate is larger than the overall death rate. In other words, the whole cell population is expected to increase infinitely. Although the total population grows infinitely, the fraction of each cell type in the population could reach a steady state. To address this question via modeling, one can rewrite Eq.(19) in terms of the fraction of each cell, i.e.,

$$f_i := \frac{X_i}{\sum_j X_j} \quad (22)$$

Hence,

$$\begin{aligned} \frac{df_i}{dt} &= \frac{X_i'}{\sum_j S_j} - \frac{X_i \sum_j X_j'}{\left(\sum_j X_j\right)^2} \\ &= \frac{\sum_j P_{ij} X_j}{\sum_j X_j} - \frac{X_i}{\sum_j X_j} \frac{\sum_j \left(\sum_k P_{kj}\right) X_j}{\sum_j X_j} \\ &= \sum_j P_{ij} f_j - f_i \sum_j \left(\sum_k P_{kj}\right) f_j \end{aligned} \quad (23)$$

Eq.(23) can be rewritten in vector format as

$$\frac{d\mathbf{f}}{dt} = \mathbf{P} \cdot \mathbf{f} - C\mathbf{f} \quad (24)$$

where $\mathbf{f} = (f_1, f_2, \ldots, f_N)$ and $C(t) = \sum_j \left(\sum_k P_{kj}\right) f_j(t)$.

Because $C(t)$ is a scalar function of time, at the steady state of Eq.(24), $C(t)$ approaches a constant, i.e., $C(t) \xrightarrow{t\to\infty} C^\infty$. In other words, the steady state of Eq.(24) is found when

$$\mathbf{P} \cdot \mathbf{f} = C^\infty \mathbf{f} \quad (25)$$

Hence, the steady state solution of Eq.(24) is an eigenvector of the coefficient matrix, **P**, normalized by the constraint, $\sum_i f_i = 1$. $C^\infty$ equals the corresponding eigenvalue of **P**. We show in the following that $C^\infty$ is in fact the largest eigenvalue of **P**.

**Theorem S1:** *The steady state solution of Eq.*(24) *is given by the normalized eigenvector associated with the largest eigenvalue of the coefficient matrix,* **P**, *with the normalization condition,* $\sum_i f_i = 1$.

***Heuristic proof:***

At $t \to \infty$, the solution to Eq.(24) approaches the solution to Eq.(26).

$$\frac{d\mathbf{g}}{dt} = \mathbf{P} \cdot \mathbf{g} - C^{\infty}\mathbf{g} \tag{26}$$

The solution of Eq.(26) reads

$$g_i(t) = \sum_k q_{ik} \exp(\lambda_k t) \tag{27}$$

where $\lambda_k$'s are eigenvalues of the matrix $\mathbf{Q} = \mathbf{P} - C^{\infty}\mathbf{I}$, and $\mathbf{I}$ is the identity matrix.

At $t \to \infty$, Eq.(27) is dominated by the term with the largest eigenvalue, i.e.,

$$g_i(t) \xrightarrow{t\to\infty} q_{i0} \exp(\lambda_{\max} t) \tag{28}$$

A nonzero steady state solution requires $\lambda_{\max} = 0$. Note that the eigenvalues of **P** have one-to-one correspondence with the eigenvalues of **Q**. For each eigenvalue of **Q**, $\lambda_k$, $\lambda_k + C^{\infty}$ is an eigenvalue of **P**. Because the largest eigenvalue of **Q** is 0, the largest eigenvalue of **P** is $C^{\infty}$.

The normalization constraint follows from the definition of fractions in Eq.(22).

**Table S1: Model parameters for DLD-1 and RPE-1 p53$^{-/-}$ cells.**

| Symbols | Meaning | DLD-1 | RPE-1 p53$^{-/-}$ | Range for data fitting (both models) |
|---|---|---|---|---|
| $b_{C2}$ | Proliferation rate of $C_2$ and SC cells | 1.2 $d^{-1}$ | 0.94 $d^{-1}$ | 0.8~1.2 $d^{-1}$ |
| $b_{C4}$ | Proliferation rate of $C_4$ cells | 1 $d^{-1}$ | 0.6 $d^{-1}$ | 0.6~1 $d^{-1}$ |
| $q$ | Probability of bipolar division in $C_2$ cells | 0.975 | 0.975 | 0.975~1 |
| $p$ | Probability that a $C_4$ cell undergoes bipolar division | 0.33 | 0.25 | Fixed * |
| $r$ | Probability of symmetric division in a bipolar division of $C_4$ cell | 0.5 | 0.7 | Fixed * |
| $fs$ | Probability that a $C_4$ cell deriving from multipolar division of $C_4$ survives | 0.4 | 0.7 | Fixed * |
| $d_{C2}$ | Death rate of $C_2$ and SC cells | 0 | 0 | Fixed * |
| $d_{C4}$ | Death rate of $C_4$ cells | 0.5 $d^{-1}$ | 0.12 $d^{-1}$ | Fixed * |
| $d_{C6}$ | Death rate of $C_6$ cells | 1.5 $d^{-1}$ | 1.5 $d^{-1}$ | Fixed * |
| $v$ | Probability of getting SC cell from a cytokinesis failure event | 0.22 | 0.32 | 0~0.6 † |
| $r_S$ | Probability that an SC cell divides symmetrically | 0.93 | 0.90 | 0.5~1 † |

† Parameters that only apply to Model II.

* Fixed at values observed or inferred from experiments.

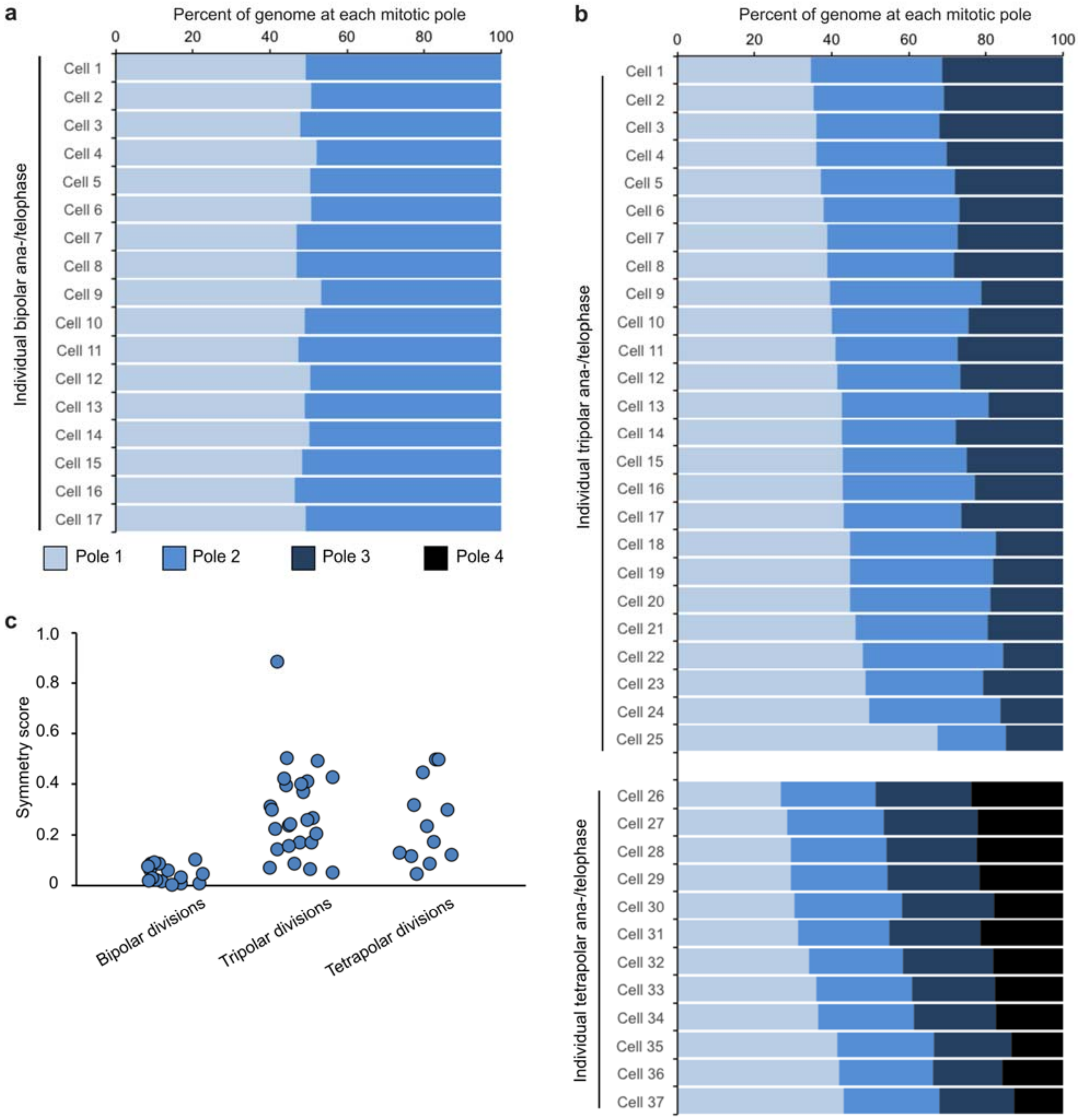


**Figure S1. DNA can be distributed unevenly to the three (tripolar) or four (tetrapolar) poles of multipolar divisions.** (a) Quantification of DAPI fluorescence intensity at the poles of individual bipolar divisions. (b) Quantification of DAPI fluorescence intensity at the poles of tripolar (top) and tetrapolar (bottom) divisions. (c) A symmetry score was assigned to each division analyzed (see methods for details). A symmetry score of zero would indicate perfect symmetry, whereas scores greater than zero indicate uneven DNA distribution among the poles.

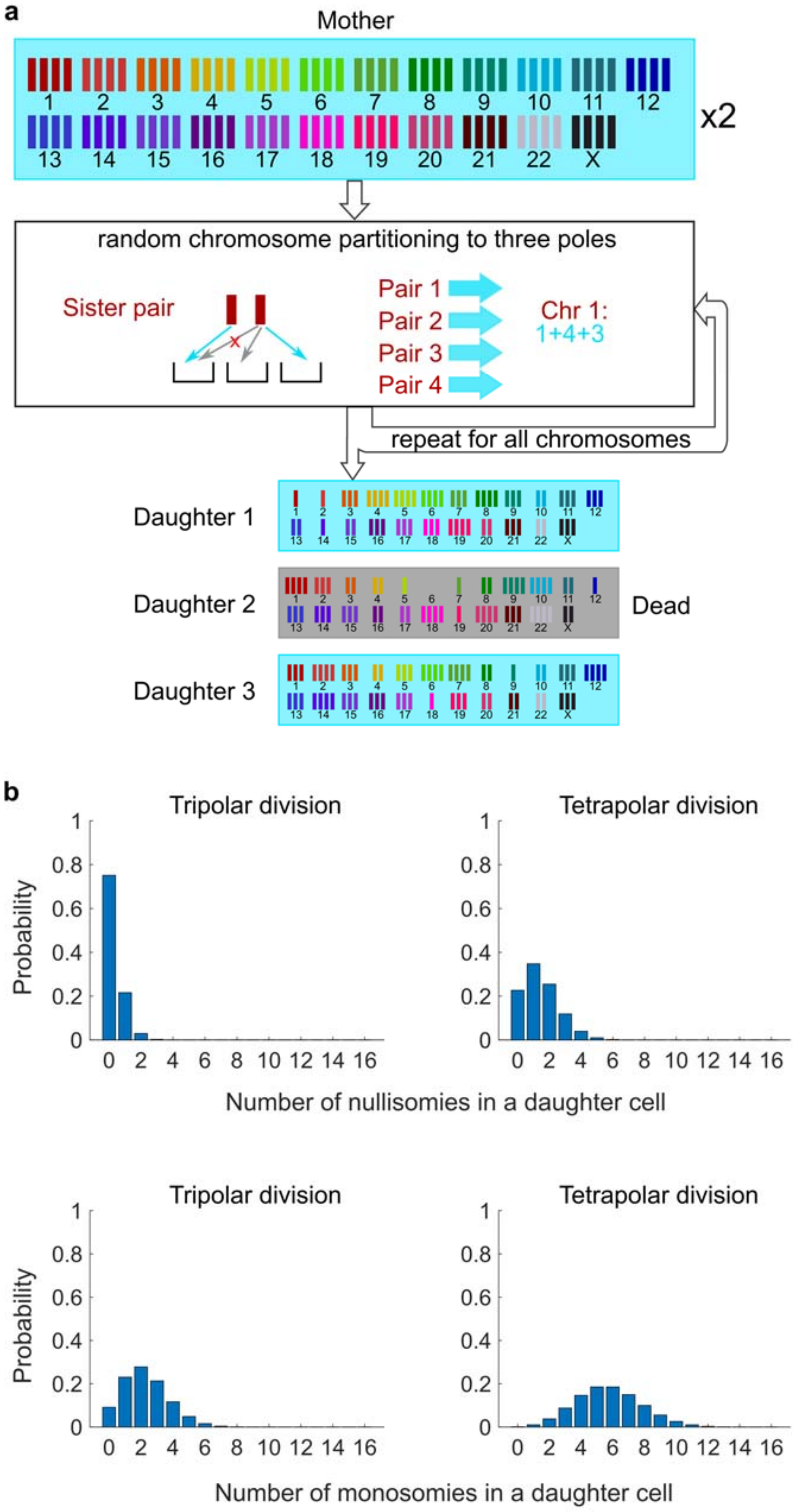


**Figure S2. Probabilistic model for karyotypic outcomes of multipolar divisions.** (a) Example result of random partitioning of chromosomes in a tripolar division. Chromosomes are randomly partitioned to three poles (cyan small arrows), with sister chromatids to different poles (red cross eliminating the case with two sisters to the same pole). Daughter 2 is dead due to nullisomy of Chromosome 6. (b) Probability distribution of the number of nullisomies (top) or monosomies (bottom) in tripolar (left) and tetrapolar (right) divisions. Calculation of the probabilities is given in Sections 1.1 and 1.2 of the Modeling Methods.

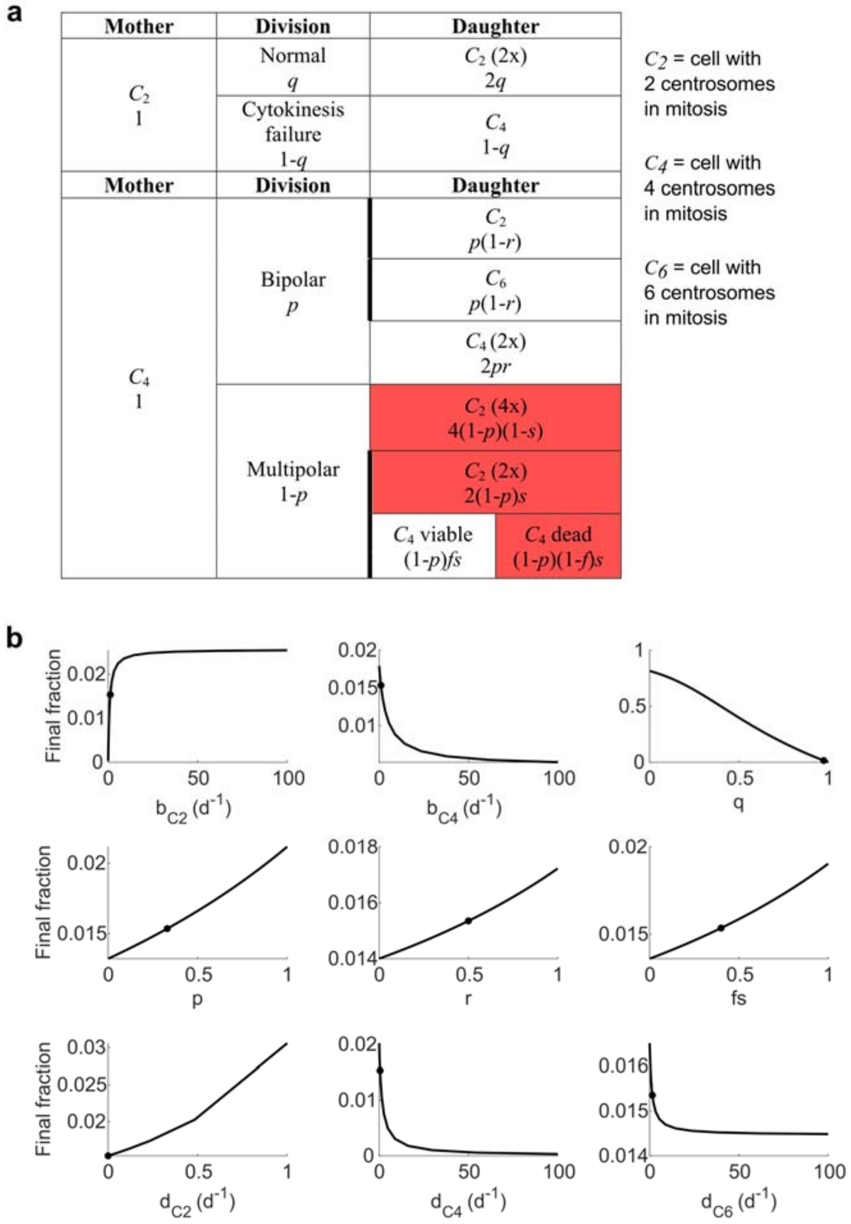


**Figure S3. Scheme and parameter sensitivity analysis for Model I.** (a) Fate of daughter cells from different types of mother cells and cell division types in Model I. Red shade: dead daughter cells. Thick borderlines: cells resulting from the same division. (b) Parameter sensitivity around nominal parameter values in Model I. Vertical axis: final steady state fraction of cells with extra centrosomes. Dot: nominal parameter for DLD-1 cells (Table S1).

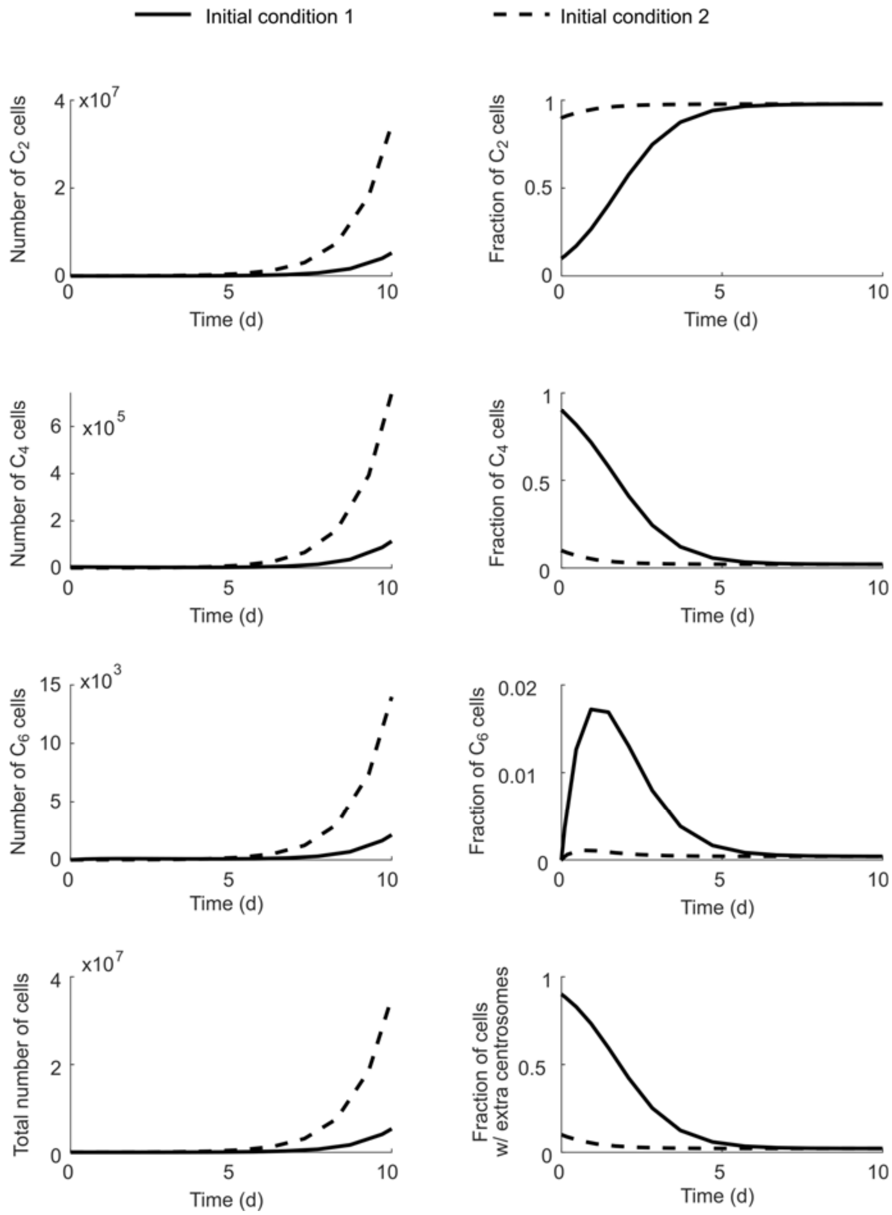


**Figure S4. Fractions of each cell type in the total population approach steady state even though the total population size grows infinitely.** Initial condition 1: $C_2(0) = 0.1$; $C_4(0) = 0.9$; $C_6(0) = 0$. Initial condition 2: $C_2(0) = 0.9$; $C_4(0) = 0.1$; $C_6(0) = 0$. Simulation results with Model I.

**a**

| Mother | Division | Daughter |
|---|---|---|
| $C_2$<br>1 | Normal<br>$q$ | $C_2$ (2x)<br>$2q$ |
| | Cytokinesis failure<br>$1\text{-}q$ | $C_4$<br>$(1\text{-}q)(1\text{-}v)$ |
| | | $SC$<br>$(1\text{-}q)v$ |
| **Mother** | **Division** | **Daughter** |
| $C_4$<br>1 | Bipolar<br>$p$ | $C_2$<br>$p(1\text{-}r)$ |
| | | $C_6$<br>$p(1\text{-}r)$ |
| | | $C_4$ (2x)<br>$2pr$ |
| | Multipolar<br>$1\text{-}p$ | $C_2$ (4x)<br>$4(1\text{-}p)(1\text{-}s)$ |
| | | $C_2$ (2x)<br>$2(1\text{-}p)s$ |
| | | $C_4$ viable $(1\text{-}p)fs$ / $C_4$ dead $(1\text{-}p)(1\text{-}f)s$ |
| **Mother** | **Division** | **Daughter** |
| $SC$ | Bipolar sym.<br>$r_S$ | $SC$<br>$2\ r_S$ |
| | Bipolar asym.<br>$1\text{-}\ r_S$ | $C_2$<br>$1\text{-}\ r_S$ |
| | | $C_6$<br>$1\text{-}\ r_S$ |

$C_2$ = cell with 2 centrosomes in mitosis

$C_4$ = cell with 4 centrosomes in mitosis

$C_6$ = cell with 6 centrosomes in mitosis

$SC$ = "super clustering" cell that efficiently clusters extra centrosomes and regularly undergoes bipolar division

**b**

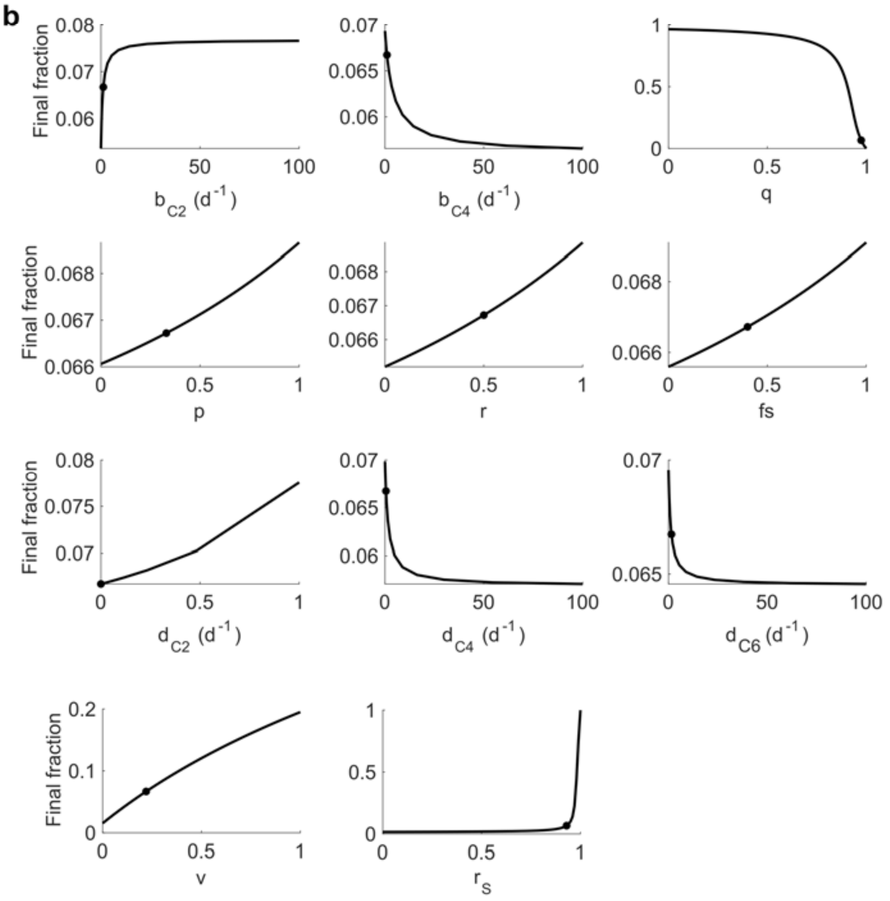


**Figure S5. Scheme and parameter sensitivity analysis for Model II.**
(a) Fate of daughter cells from different types of mother cells and cell division types in Model II (with SC cells; i.e., $C_4$ cells that can effectively cluster extra centrosomes, and regularly undergo bipolar division). Red shade: dead daughter cells. Thick borderlines: cells resulting from the same division.
(b) Parameter sensitivity around nominal parameter values in Model II. Vertical axis: final steady state fraction of cells with extra centrosomes. Dot: nominal parameter for DLD-1 cells (**Table S1**).

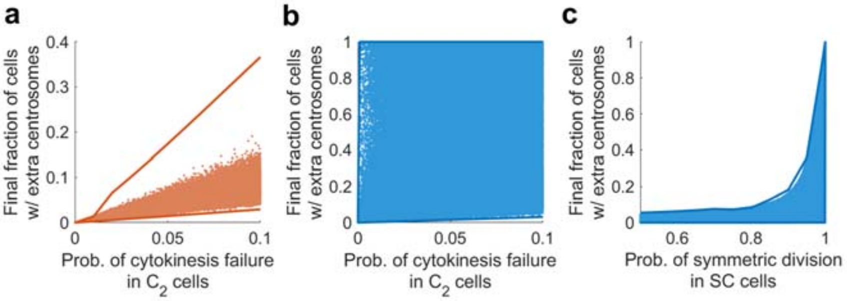


**Figure S6. Final fractions of cells with extra centrosomes are strongly constrained by the most sensitive parameters in both models.** Solid lines: upper and lower limits of the final fraction computed through optimization. Scattered dots: final fractions computed over 1 million randomly sampled parameter sets. (a) In Model I the final fraction of cells with extra centrosomes is strongly constrained by the probability of cytokinesis failure in $C_2$ cells. Note that the upper limit stays far away from the common range of results from random parameter sets. This phenomenon indicates that the upper limit is approached via very stringent conditions on the parameter values, and is likely not robust. The range covered by the scattered dots are robust and hence more realistic. (b) In Model II, the final fraction of cells with extra centrosomes is not constrained by the probability of cytokinesis failure in $C_2$ cells. (c) In Model II, the final fraction of cells with extra centrosome is constrained by the probability of asymmetric division in SC cells. In (a-c), the final (i.e., steady state) fraction of cells with extra centrosomes were computed using the method posited in Theorem S1. Distributions of random parameter sets chosen within physiologically reasonable range: $b_{C2}$ ~ log uniform [0.8, 1.2], $b_{C4}$ ~ log uniform [0.1, 1.5], $q$ ~ uniform [0.9, 1], $p$ ~ uniform [0. 0.6], $r$ ~ uniform [0, 0.9], $fs$ ~ uniform [0, 0.9], $d_{C2}$ ~ log uniform [$10^{-10}$, 0.3], $d_{C4}$ ~ log uniform [$10^{-10}$, 1], $d_{C6}$ ~ log uniform [0.5, 2], $v$ ~ uniform [0. 0.6], $r_S$ ~ uniform [0.5, 1].
